# High-Resolution Spatial Transcriptomics Reveals Millimeter-Scale Malignant Territories and Immune Exclusion patterns in Human Salivary Gland Adenoid Cystic Carcinoma

**DOI:** 10.64898/2026.11.10.765744

**Authors:** Gopikrishnan Bijukumar, Kendall Hoff, Kathryn J. Brayer, Elaine L. Bearer, Viswanathan Palanisamy, Jeremy S. Edwards

## Abstract

**Background/Objectives:** Adenoid cystic carcinoma (ACC) of the salivary gland (SGACC) is a rare malignancy with frequent late recurrence and limited response to systemic and immune therapies. Bulk and single-cell sequencing do not preserve tissue architecture, so the spatial organization of its MYB-driven myoepithelial and luminal tumor cells, stroma, and immune cells remains poorly defined. This study aimed to map the SGACC tumor microenvironment at near-cellular resolution.

**Methods:** Visium HD based spatial transcriptomics was performed on four SGACC tissue sections from three patients with tubulo-cribriform tumor form in stage III - IV. Bins of 10 x 10 µm were annotated and deconvolved with cell2location using an integrated single-cell reference of 119,189 cells from 20 donors. Pseudobulk differential expression, spatial autocorrelation, potency and spatial trajectory inference, permutation-based territory analysis, Hotspot module detection, and marker-based immune profiling were applied.

**Results:** MYB-expressing luminal (LECs-MYB) and myoepithelial (MECs-MYB) epithelial cells occupied 55.7% of bins on average. The MECs-MYB to LECs-MYB ratio increased from the nest interior toward the tumor edge, where cancer-associated fibroblasts rose sharply. Potency and trajectory analyses placed MECs-MYB upstream of LECs-MYB. Malignant states formed territories of up to 38.0 mm² that persisted over 1.5 mm in three sections and contained spatially restricted programs. Hypoxic stressed territories also showed mTORC1, unfolded protein response, and cell-cycle programs. T/NK, myeloid, MHC class I and II, and plasma cell signatures were 1.4-to 5.7-fold higher in stroma than in tumor nests, whereas VTCN1 (B7-H4) was the only B7 family ligand enriched within nests.

**Conclusions:** SGACC shows an ordered arrangement of malignant lineages, millimeter-scale malignant territories, MHC class – I and II enriched stroma, and immune-excluded nests enriched for B7-H4. These features provide evidence to be validated in larger cohorts with protein-level and clinical data.

**Simple Summary:** Adenoid cystic carcinoma is a rare cancer of the salivary glands that often returns years after treatment and responds poorly to current drugs and immunotherapy. Most earlier studies broke the tumor apart before measuring gene activity, losing the localization of cell in tumor. In this study, we measured gene activity directly on four thin tissue slices from the tumors of three patients at near single-cell detail. The two main types of cancer cells were arranged in an ordered pattern within tumor clusters, and cancer cells of the same type formed large regions several square millimeters in size, each with its own local gene activity. Immune cell signals stayed mostly in the surrounding supportive tissue and rarely reached the tumor clusters, while the tumor cells showed high activity of the gene for B7-H4 that can dampen immune responses. The patient with the longest recorded survival had an altered territory make up from that of other patients that points towards features that studies with larger cohort size can test and develop targeted treatment approaches.

## Introduction

Adenoid cystic carcinoma (ACC) is one of the most clinically distinct, rare, malignant neoplasms of salivary glands in humans. The annual incidence is 3-4.5 cases per million worldwide. ACC accounts for almost 10-25% of salivary gland cancers and contributes to 1% of all head and neck malignancies. ACC arises in both minor and major salivary glands with a preference to regions comprising luminal and secretory epithelium[1–3]. Despite its rarity ACC remains a major clinical challenge because of frequent late recurrence and limited therapeutic options available till date. Clinically, ACC is characterized by its indolent but aggressive nature over time. In addition to local recurrence, ACC exhibits a tendency for delayed distant metastasis, most commonly to the lungs but also to bone and liver in ∼30-40% of patients[2,3]. A feature of ACC that is frequently associated with poor prognosis is perineural invasion which contributes to deep local extension near neurons that makes its complete surgical resection nearly impossible[4]. ACC is also notable for its biphasic cellular composition, primarily containing the epithelial and myoepithelial cells that make up its characteristic morphology and phenotypic complexity[2]. Histopathologically, ACC is frequently divided into 3 categories, tubular, cribriform and solid tumor morphology[1]. Interestingly these tumor growth patterns are also associated with patient prognosis. The tubular subtype is frequently associated with favorable prognosis whereas the cribriform subtype is associated with intermediate prognosis[2,3]. Solid subtype is categorized as the most aggressive and is associated with higher grade disease, greater metastatic potential and poor prognosis[2].

At the molecular level, advancements in next-generation sequencing (NGS) technologies have provided significant insights into the gene-regulatory networks of this malignancy[2,5]. The well-regarded genetic hallmark is the t(6;9) chromosomal translocation that leads to MYB - NFIB fusion which is seen in 50-78% of cases[4]. These chromosomal events lead to the activation of the transcription factor MYB that leads to un-controlled cell cycle progression, resistance to apoptosis, upregulation of angiogenesis and epithelial to mesenchymal transition related programs[2]. Beyond MYB alterations, recurrent mutations in the NOTCH pathway including NOTCH1-4 have also been documented as important contributors to ACC[5]. Additionally, recent studies have also drawn correlations in chromatin remodeling genes to ACC such as SMARCA2, KDM6A, ARID1A[6]. Recent proteogenomic studies have taken a big step by defining two molecular subtypes of salivary gland ACC (SGACC) called ACC - I and ACC - II. ACC - I is characterized by the strong upregulation of MYC and enrichment of NOTCH1 alterations that leads to a more clinically aggressive behavior whereas ACC - II shows upregulation of TP63 and receptor tyrosine kinases such as AXL, MET and EGFR and is associated with a more indolent clinical course[7].

The treatment modalities used to treat ACC has historically remained limited. Surgical resection with post-operative radiotherapy continues to be the cornerstone for ACC tumors. These treatment methods do not focus on limiting the local recurrence and metastatic potential of this malignancy. Conventional cytotoxic chemotherapy has shown modest benefits, while targeted therapies and immune checkpoint target and blockading methods have yielded limited clinical responses[8]. ACC shows this resistance because of its distinctive tumor microenvironment, which is typically ‘immune cold’ with low PD-L1 expression and very low tumor infiltrating lymphocytes[9,10]. Spatial immunoprobing has revealed that even tumors with relatively higher immune cell content often show stromal separation of immune cells instead of effective infiltration into the tumor[11].

Despite significant advances in bulk transcriptomics, single cell transcriptomics and genomic studies there are huge gaps in our understanding of ACC biology. Bulk RNA seq (used most often in ACC studies) provides an average transcriptional makeup of ACC tumor while obscuring regional and cellular variation. Single cell RNA seq preserves the cellular resolution that facilitates the understanding of cells that make up the tumor microenvironment (TME) but it disrupts the tissue, losing the spatial information that helps to determine the physical proximity of cells. For a deeper understanding of the complex makeup of the ACC TME, we need to incorporate spatial transcriptomics-based approach as it can capture high resolution transcriptional states across preserved tissue topology, enabling the understanding of tumor heterogeneity, cellular neighborhoods and microenvironmental niches[11].

In this study we are applying Visium HD based spatial transcriptomics approach to a cohort of 4 human SGACC tumor sections from 3 patients **(Table. S1)** to generate a high-resolution spatially resolved view of this tumor associated to complex malignancy. This approach will help us map the tumor microenvironment, define spatial organizations of malignant and stromal compartments and characterize the immune infiltration patterns. Particularly the MYB vs non-MYB expressing cell populations existing within the tumor will be mapped and studied to identify the molecular differences and spatially restricted gene expression programs. The MYB expressing cell types will be closely studied relative to each other, together with whether these malignant states are organized into territories or intermixed within the TME. This study will also focus on unraveling TME domains and their associated functions. Through this approach we aim to deepen the understanding of ACC biology by providing a high-resolution view of the gene expression programs and spatial organization within the complex SGACC TME.

## Materials and Methods

1. **Binning / pre-processing / clustering** To optimize the signal-to-noise ratio and reach a single cell resolution, the raw data originally comprising 25 million features at a 2 x 2 µm resolution underwent a two-step spatial binning process. First, features were binned into 10 x 10 µm units by aggregating 5 x 5 features along X and Y spatial axes. Next a 5 x 5 sliding window convolution was applied for further smoothing the spatial expression profiles. The 10 x 10 µm resolution was used for downstream clustering and annotations. The quality control and filtering was done in python using Scanpy (v1.11.5)[12]. Quality control metrics such as total transcript counts and number of genes detected per bin was estimated using *sc.pp.calculate_qc_metrics*. Bins with a total transcript and gene counts lower than thresholds determined for each sample from QC plots were excluded. This helped in removing sparsely captured bins prior to downstream analysis. The data was then normalized using *sc.pp.normalize_total* which was followed by log1p transformation via sc.pp.log1p function. Finally, the QC data was re-calculated to confirm the effects of filtering resulting in clean data fit for downstream analysis. The data from 4 sections were concatenated into a single Anndata object[13] using the *anndata.concat* function. The highly variable genes (HVGs) were selected using the Seurat v3[14] variance-stabilization approach in the *sc.pp.highly_variable_genes* function. Top 5000 HVGs were chosen and subsetted. To see how strongly the samples separated before any correction, we clustered the data without batch correction. Rapids-singlecell (v0.13.4)[15] package in python was used for accelerating this step on GPU. Next, the top 30 principal components were computed followed by neighborhood graph generation with k = 30. This was clustered using the Leiden clustering at a resolution of 0.5. Random seed was set to 42 for every stochastic step.
2. **Batch correction using scVI** Batch effects were corrected using single-cell variational inference (scVI) with sample ID **(Table. S1)** set as the categorical covariate. The model was set up using *scvi.model.SCVI.setup_anndata* and the raw counts were provided as input. The model was trained on raw UMI counts of the 5,000 highly variable genes with a negative binomial likelihood. The encoder mapped each bin to a 30-dimensional latent space, and a decoder reconstructed the counts from this latent representation. Training was GPU accelerated with a batch size of 8192 and a fixed learning rate of 1e-03, with a maximum of 400 training epochs. Early stopping was set up with a patience level of 20 epochs, so that the training terminated when the improvement in the validation curve stopped. The training and ELBO validation was plotted and examined to confirm convergence. Once the model converged the batch corrected latent representation was extracted using *model.get_latent_representation* and stored in the adata.obsm attribute of the Anndata object. The neighborhood graph was recomputed in the new latent space and clustered using Leiden algorithm (v0.10.2) at a resolution of 0.5. All downstream cell type annotations and spatial analyses were conducted on this scVI batch corrected embedding. Finally, the samples were split based on batch information for independent downstream analyses. The 4 samples were split into their respective H5ADs. After inspecting the UMAPs of individual samples, it was concluded that the UMAP structure needs to be recomputed to unravel the transcriptional states in each sample. The steps were accelerated on GPU using the rapids-singlecell package. The k-nearest neighbor graph was recomputed in scVI latent space with k set to 50 for all the samples for un-biased comparisons. This was followed by dimensionality reduction using UMAP with a *min_dist* parameter set to 0.6. To identify the optimal clustering resolution the Leiden algorithm resolution was varied ranging from 0.1 to 1.0, and normalized mutual information(NMI) were plotted. The resolution contributing to highest NMI value was chosen for cluster based manual annotation and downstream analysis.
3. **Single cell Reference dataset preparation** This SGACC spatial dataset was not complemented by a single cell dataset. To address this matter, we made a reference by combining 5 single cell publicly available datasets. These included the GEO datasets from GSE210171, GSE342688, GSE217084, GSE294017 and the Tabula Sapiens salivary gland atlas **(Table. S2)**. This ensured a representation of malignant and normal cell types in ACC and human salivary gland. After pre-processing and filtration, raw counts were integrated with scVI (30 latent dimensions) on 4000 highly variable genes with batch (library or patient) and technology as covariates. Malignant cells were identified by the expression of ACC specific genes such as MYB, MYBL1, EN1, SOX11 and FABP7 compared cluster by cluster with the normal salivary gland epithelium along with copy number variation profiles from inferCNV (v0.6.1) as supporting data. The cell types were then annotated within each major compartment (epithelial, stromal, endothelial and immune) using canonical marker genes, the expert labels in of Tabula sapiens and CellTypist (v1.7.1)[16], and confirmed by pseudobulked differential expression analysis. Finally, the reference contained 119,189 cells from 20 donors, including 32,009 malignant cells grouped into 34 cell types. This reference single cell dataset was used to perform reference-based deconvolution.
4. **Deconvolution using Cell2Location**

a. **Reference Signature Estimation** Cell type expression signatures were estimated from the integrated single-cell reference dataset with a negative binomial regression model of cell2Location (v0.1.5, scVI tools v1.3.3). The model was fitted on raw counts from 118,820 cells (after preliminary filtering) across 34 different cell types. The sample ID was used as the batch variable and sequencing technology (10x Genomics single-cell, 10x Genomics single-nucleus or Smart-seq2) as additional categorical covariates. The genes in the reference single-cell dataset that were also present in our spatial samples were only considered for model fitting. The mitochondrial genes were dropped and genes with low expression were removed during cell2location filter / preparation step (*cell_count_cutoff* = 25, *cell_percentage_cutoff2* = 0.1, *nonz_mean_cutoff* = 1.2), leaving 18,508 genes. The model was trained on all cells for 400 epochs (minibatch size 2,500, learning rate 0.002). All the samples had the ELBO loss plot plateaued. The cell type signature was the posterior mean expression of every gene from the last 1500 posterior samples.
b. **Spatial Abundance Mapping** The cell type abundances of each of the 4 spatial samples were separately mapped with the cell2location spatial model using the same reference signatures. There were almost 4,250 genes that were common between the reference signatures and the spatially variable genes from the tissue. The expected number of cells per bin (*N_cells_per_location*) was set to 3 to reflect the small size of Visium HD bins, and *detection_alpha* was set to 20, the recommended value for regularizing within-section variation in detection sensitivity. All bins were used for training (*train_size* = 1) in minibatches of 10,000. A ClippedAdam based optimization was used with an initial learning rate of 0.002 and a gradient norm clip of 10. The learning rate decayed exponentially to 1% of its starting value by the final step to stabilize the training. Parameters from the epoch with the lowest training loss were retained. Cell abundance per bin was summarized from 1,000 posterior samples. These abundances were used for spatially restricted enrichment of cell types on the tissue.
5. **Pseudobulked differential expression analysis with PyDESeq2** We compared the gene expression patterns between tissue regions with PyDESeq2 (v0.5.4) using pseudobulk profiles from the four ACC tissue sections. The bins were taken in 50 x 50 µm window spaced 10 µm apart. In this configuration the expression patterns of neighboring bins always spill over affecting the analysis. Hence, we kept only the bins at every fifth grid position in both directions, which gave non-overlapping 50 × 50 µm tiles in which each molecule is counted once. Tiles labeled as low quality were removed. Raw counts of the remaining tiles were summed up by section and region (tumor nests, stroma and normal gland). The gene that were present across all 4 samples with at least 10 transcripts were only considered for analysis. As stroma is usually intermixed with the tumor nests in SGACC tissues, for the analysis only the core regions of stroma and tumor nests were used. A Euclidean distance transform on the full bin grid gave, for each tumor bin, the distance to the nearest stromal bin and, for each stromal bin, the distance to the nearest tumor bin. Bins at least 80 µm from the other compartment were defined as core, and their tiles were summed into one tumor-core and one stroma-core pseudobulk per section. For each run the contrast parameter of PyDESeq2 was set with the design ∼ section + region. P values came from Wald test, and multiple testing correction was done using Benjamini-Hochberg method[17]. Shrunk log2FC was used with apeGLM and genes with an adjusted P (FDR) < 0.05 and | shrunken log2FC | ≥ 0.5 were considered significant. The setup was in such a way that positive values mean, enriched in tumor nests. Gene set enrichment analysis was performed using GSEAPy (V1.1.11). All tested genes were ranked by the Wald statistic without filtering and was run against the MSigDB Hallmark, Reactome and GO Biological Process collections, keeping gene sets of 15 to 500 genes. Each run used 1000 permutations and fixed random seed. Gene sets with an FDR < 0.05 were considered significant, and a positive normalized enrichment score (NES) indicated enrichment in tumor nests.
6. **Spatial auto-correlation analysis with Moran’s I** We computed Moran’s I for every gene across all 4 tissue sections using the same non-overlapping 50 µm tiles used in the previous analysis. Raw counts of 17,943 genes present in all four tumor sections were normalized to the median tile total and log1p transformed. A spatial graph was built in Squidpy (v1.8.1) linking each tile to its eight adjacent tiles. Tiles without a neighbor were dropped. Genes detected in at least 2% of tiles and in at least 20 tiles were used for downstream steps. Some genes show a technical artifact in which their counts were concentrated along a single row or column of the capture area. Moran’s I, might consider this as a false positive by considering it as a spatial cluster. We flagged a gene as a stripe artifact when two of the conditions were met. A single tile row or column had at least 15% of its counts. Second, Moran’s I differed by at least 0.1 between a calculation using only left-right neighbors and one using only up-down neighbors, with the higher value along the direction of the stripe. A gene was called spatially variable when its FDR is less than 0.05 and I is greater than or equal to 0.1 and didn’t fall into stripe artifact. Top 300 genes with highest I value was used for identifying spatial gene co-expression modules. Expression of these genes was z-scored and smoothed by averaging each tile with its neighbors. A gene-gene Pearson correlation (r) was computed on these smoothed values. Genes were clustered by average linkage on 1 − r and the tree was cut at 0.6 and clusters with at least 10 genes were kept as modules. The modules whose scores correlated with the log tile depth (|r| ≥ 0.5) were flagged as possibly driven by sequencing depth. The identified modules from different spatial samples were compared pairwise with a one-sided hypergeometric test. The genes tested in both samples were used as the universe, and Benjamini-Hochberg corrections were applied to all tests. The modules that showed similarity with FDR < 0.05 were considered to be modules performing similar functions across tumor samples, defined as groups. The gene networks were drawn in python using igraph (v1.0.0)[18].
7. **Spatial trajectory analysis using SpaTrack** Bins in the spatial dataset are noisy. Hence, the root of the trajectory was set by comparing the developmental potential of MECs-MYB and LECs-MYB with CytoTRACE2 (v1.1.0)[19], in five tumors of one 10x Genomics 3’ dataset of the single cell reference (GSE342688; SC1 to SC5). CytoTRACE2 scores scale with the number of detected genes so every cell was down sampled to 4,728 UMIs before scoring. The lineage difference in potency score was tested with a linear mixed model (lme4 (v2.0.1)) with S and G2M cell-cycle scores as covariates. Sensitivity analyses were done by dropping cell-cycle terms and through leave one sample out tests. Monocle3 (v1.4.27) was run on the depth matched counts with the graph rooted in the clusters with at least 90% MECs-MYB (through CytoTRACE2 potency scoring). For the spatial analysis, we used the non-overlapping 50 µm bins. Each sample was separately analyzed with spaTrack (v1.0.2) after regressing out depth-based variations. SpaTrack links bins by entropy-regularized optimal transport. In our analysis we weighted expression and physical distance equally and ranked each bin by the transport mass it receives from a set of start bins. The start was 50 MECs-MYB bins from five k-means clusters of their positions; ACC3, which has no MECs-MYB, was started from MEC-like stressed bins. Lineage separation was summarized as the AUC of LEC-like (LECs-MYB and LEC-like stressed) versus MEC-like (MECs-MYB and MEC-like stressed) bin pseudotime, overall and within read-depth quintiles. For pathway analysis the raw counts from tumor bins in each 1/3^rd^ of the pseudotime trajectory was summed in each section. Genes were ranked using the Wald statistic and tested for Hallmark and Reactome enrichment with fgsea (v1.32.4) and Benjamini-Hochberg correction. The genes that were detected in at least 1% of reference MECs-MYB or LECs-MYB were ranked, which excluded background signal for cleaner enrichment patterns from spatial data.
8. **Spatial organization of tumor states and characterization of stressed territories** To test whether the three tumor states were intermixed or grouped into territories, we examined every pair of tumor bins up to 1.5mm apart. For each state and distance, we calculated how often a bin of that state had a neighbor of the same state and divided this by the share of that state among all tumor bins. This co-occurrence ratio equals 1 when the states are randomly mixed. To show what random mixing looks like, we shuffled the state labels among the tumor bins of each section 1,000 times. The shuffles keep the shape of the nests and the number of bins of each state, and they provided the P values and 95% bands for the co-occurrence ratio. As a state can also look clustered simply because it fills a nest of its own, we repeated the shuffle analysis within each tumor nest. Touching bins of the same state, including diagonal neighbors, were joined into territories with scikit-image (v0.26.0). For each state, we calculated the percentage of its bins that lay in territories of 1 mm² or larger. To find out what other terms are enriched besides hypoxia in the stressed territory, we compared the stressed tumor cells-MYB with MECs-MYB in ACC1, 2, and 4, and LECs-MYB in all four sections. Again, in order to avoid mixing, we only considered the core bins that lie at least 100 μm away from any bin of a different state. The raw counts of these bins were summed per sample and per cell population, and the resulting pseudobulk states were compared using PyDESEq2 (v0.5.4). The contrast was set with the design ∼ section + state. A gene was called significant when its FDR values is less than 0.05 and its absolute apeGLM-shrunken log2 fold change was at least 0.5. A differentially expressed gene was considered robust if the expression changed in the same direction across all samples and even through a leave one out analysis. Pathways were tested by pre-ranked gene set enrichment analysis on the MSigDB Hallmark gene sets with GSEAPy (v1.1.11, 1,000 permutations, FDR below 0.05). Genes were ranked by the Wald statistic. Lastly, to see whether the stressed program switches on sharply at the territory border or rises gradually, we averaged the scores and VEGFA expression in 50 μm steps of distance on both sides of the border and compared across it.
9. **Territory restricted gene expression program analysis with Hotspot** We ran Hotspot (hotspotsc v1.1.3) analysis on the largest malignant territories of MECs-MYB and LECs-MYB that we got across all the tumor tissues. This gave us seven spatially restricted gene expression modules exclusively present in the tumor nests. Within each territory, genes with a mean expression above 20 counts per million (CPM) that were detected in more than 5% of tiles were tested, excluding mitochondrial, ribosomal and non-coding genes. The number of nearest neighbors (k) was set to 28, which links every 50 µm tiles to the tiles in its surrounding 150 µm. The 500 most strongly autocorrelated genes (FDR < 0.01) were grouped by pairwise local correlation into modules of at least 10 genes, and each tile was scored for every module. Near the stressed territory, modules that overlapped the Hallmark hypoxia and increased towards the stressed border were removed as part of the stressed signal. The stability of the obtained modules was checked by repeating hotspot analysis with variable number of neighbors (15 and 50) and also by scoring the program genes in the single cell reference dataset.
10. **Immune landscape analysis at the tumor nest edge** The immune analysis used the same 50 µm tiles and the 17,943 genes shared by the four sections, and expression of a gene set in a group of tiles was given as pooled CPM (summed gene-set counts divided by summed tile depth). Each tile was given a signed distance to the tumor nest edge. The tumor cells within the nest had negative distance and every other bin outside had positive distance towards it. Immune signal was measured with marker gene sets for T/NK cells (CD3E, CD3D, CD2, CD8A, NKG7, GZMB, GZMK, CCL5), myeloid cells (CD163, C1QA, C1QB, LYZ, CD14, AIF1), B cells (MS4A1, CD79A, CD19, CR2), tertiary lymphoid structures (CXCL13, CCL19, CCL21, LAMP3), plasma cells (JCHAIN, MZB1, TNFRSF17, DERL3) and interferon response (STAT1, IRF1, CXCL9, CXCL10, GBP1, IDO1). MHC class II was represented by CD74 alone, as the HLA-D genes were absent or nearly undetected. Each gene set was compared with 100 control sets in which every gene was replaced by a random gene of the same expression decile, and its percentile among them was reported. This allowed us to determine if the immune signal is actually present or just because of the differences in the bins that we are comparing. The tumor-intrinsic genes VTCN1 (B7-H4), B2M, TAP1, PSMB8 and PSMB9 (MHC class I), CD274 (PD-L1), PDCD1LG2 (PD-L2) and CD276 (B7-H3) were compared between the nest interior of each tumor state and normal gland, before and after down sampling tiles to the 10th-percentile depth of their section. Plasma-cell isotypes were quantified by measuring each heavy-chain signal (IGHA1, IGHA2, IGHG1, IGHG2, IGHG3, IGHM, IGHD, IGHE) among all heavy-chain counts in nest, stromal and normal gland zones, and nest tiles. We only considered bins with at least five heavy chain transcript counts as we wanted to separate actually infiltrating vs potentially ambient immunoglobulin transcripts.

## Results

### Independent sample annotation reveals common cell types spanning across SGACC tumor samples

Our first approach to understand and map the SGACC TME was to investigate the cell types that make up each tumor sample. The ACC tumor tissues were processed independently on 4 different Visium HD spatial chips to perform spatially resolved analysis and spatial projection on the tumor tissue **(Fig. 1A)**. The spatially resolved gene expression data was subjected to custom binning (10 x 10 µm), standard pre-processing, and filtering steps. Unsupervised clustering using Leiden algorithm[20] showed sample-based confounding **(Fig. 1B)**. To eliminate this batch effect, single cell variational inference[21] (scVI) (v1.4.2) based batch correction was implemented **(Fig. 1C)**. The batch corrected samples were split and independently annotated, by considering highly expressed marker genes from each cluster in each tissue. A cluster was called MYB-expressing ACC tumor only if the MYB was detected in its bins more often than about 200 control genes of similar expression level (after correcting for sequencing depth), and only if its enrichment held in bins far from other tumor clusters **(Fig. 1D; Fig. S1A)**. The identified and annotated cell types were projected spatially on the tumor tissue to map out the TME of the ACC tumor tissue **(Fig. 1E)**.

**Figure 1.**
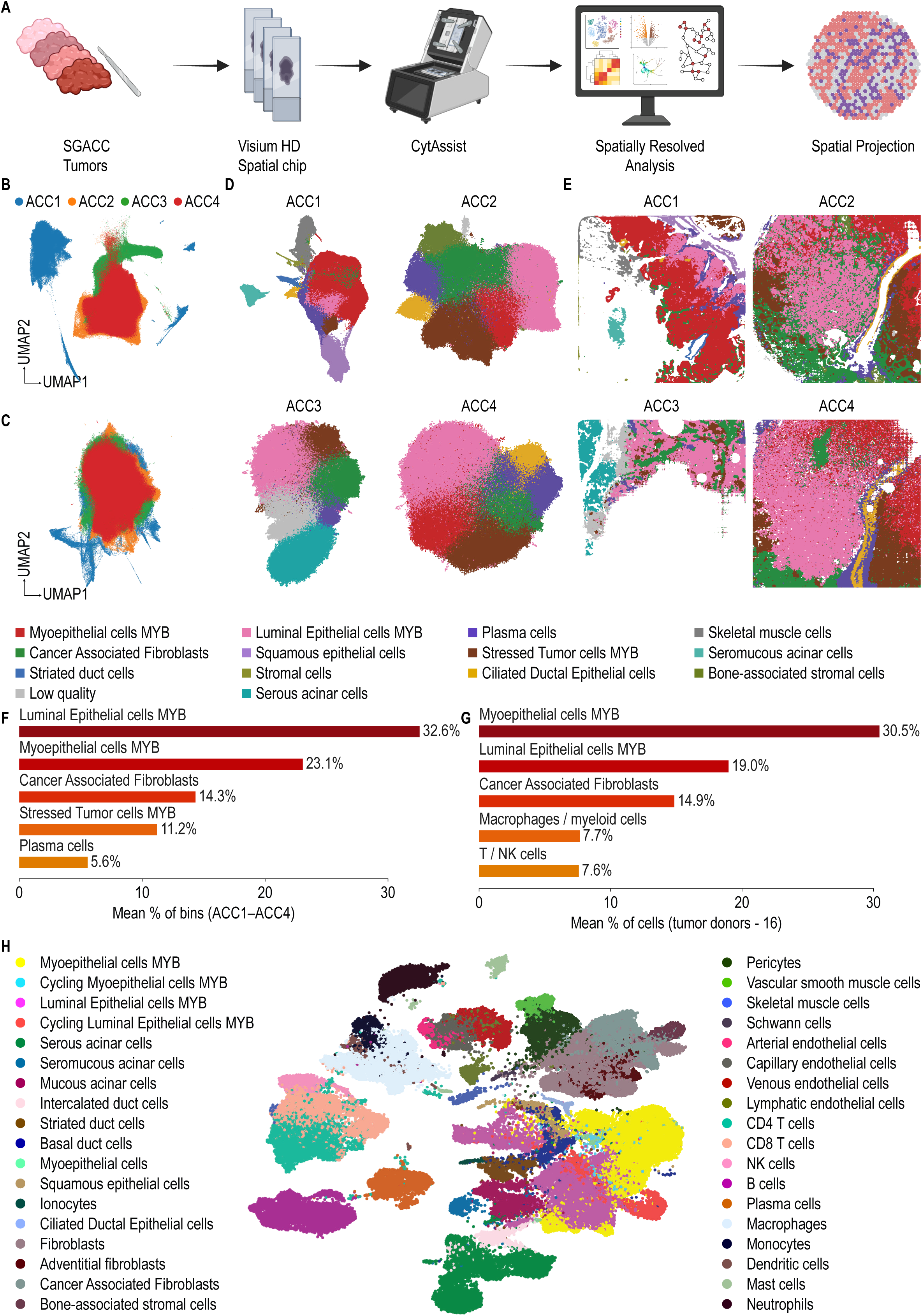
Cell-type annotation of four SGACC tumor sections and the single-cell reference. **(A)** Workflow. Four salivary gland adenoid cystic carcinoma (SGACC) tumor sections were profiled on different high resolution spatial chips. The spatially resolved. analysis was done and the results were projected back onto the tissue. **(B)** UMAP of the bins of all four sections (ACC1 to ACC4), colored by section, before batch correction. **(C)** The same bins after batch correction with single-cell variational inference (scVI). **(D)** UMAP of each tissue section colored by annotated cell type after re-clustering. **(E)** Annotated cell types mapped onto each tissue section. Colors in (D) and (E) follow the legend below (E). **(F)** The five most abundant cell types in the spatial data, given as the mean percentage of bins across ACC1 to ACC4 (low-quality bins excluded). **(G)** The five most abundant cell types in the single-cell reference, given as the mean percentage of cells across the 16 tumor donors (normal and adjacent normal samples excluded). **(H)** UMAP of the integrated single-cell reference (119,189 cells from 37 samples of 20 donors; four ACC datasets and the Tabula Sapiens normal salivary gland), colored by cell type. In the spatial data, the suffix MYB marks cell types whose clusters passed the depth-controlled MYB enrichment test.

Averaged across the four samples, luminal epithelial cells-MYB (LECs-MYB) were the most abundant cell type, accounting for 32.6% of 10 x 10 μm bins. Myoepithelial cells-MYB (MECs-MYB) ranked second at 23.1%. Together, these two populations made up more than half of all bins averaged across four ACC tumor samples (55.7%) **(Fig. 1F)**. At the individual sample level, interestingly, LECs-MYB was the most abundant cell type in three of the four samples. ACC1 was the exception, with MECs-MYB occupying a larger proportion of bins **(Fig. S1B)**. As the tumor sample set was not accompanied by a single-cell dataset, we built a comprehensive reference by combining ACC-affected salivary gland and normal salivary gland single-cell samples from five public datasets **(Table. S2)**. Four were ACC cohorts (GSE210171, GSE342688, GSE217084 and GSE294017) and the fifth was the normal salivary gland from Tabula Sapiens[22–26]. The final reference dataset contained 119,189 cells from 37 samples of 20 donors (16 with ACC, 4 healthy) **(Fig. 1H).** A total of 34 cell types were identified that could be broadly categorized into malignant, stromal, vascular and immune cell populations **(Fig. S1C)**. We identified malignant cells by their elevated MYB expression relative to normal salivary gland epithelium as MYB is a known driver of ACC **(Fig. S1D)**. Interestingly, the most abundant cell types in the tumor tissue closely matched single-cell reference dataset. Myoepithelial cells MYB, luminal epithelial cells MYB, and cancer-associated fibroblasts were the most abundant cell types in the single-cell reference dataset matching the dominant cell populations spatial dataset **(Fig. 1G)**. This suggests that the reference can support accurate estimation of cell type abundances in our tissue with cell2location[27], as the major tissue cell types are well represented in the reference.

### Cell2Location resolves the distribution and estimates the spatially confined enrichment of malignant and non-malignant cell types in SGACC TME

Cluster-based annotation of the four SGACC tumor samples assigns a single cell type identity to each 10 x 10 μm bin, so it cannot capture the cellular composition of bins that contain more than one cell type. To estimate the abundance of each cell type we performed deconvolution across 2.5 million bins across the four SGACC tumor sections with Cell2Location (v0.1.5) **(Fig. 2A)**. The cell2location regression model was trained on the single cell reference data with sample and sequencing technology as covariates to learn the cell type specific signatures. Each spatial sample was then mapped against these signatures with the cell2location hierarchical Bayesian model. Cell abundances were estimated from the epoch with the lowest training loss **(Fig. S2A)**.

**Figure 2.**
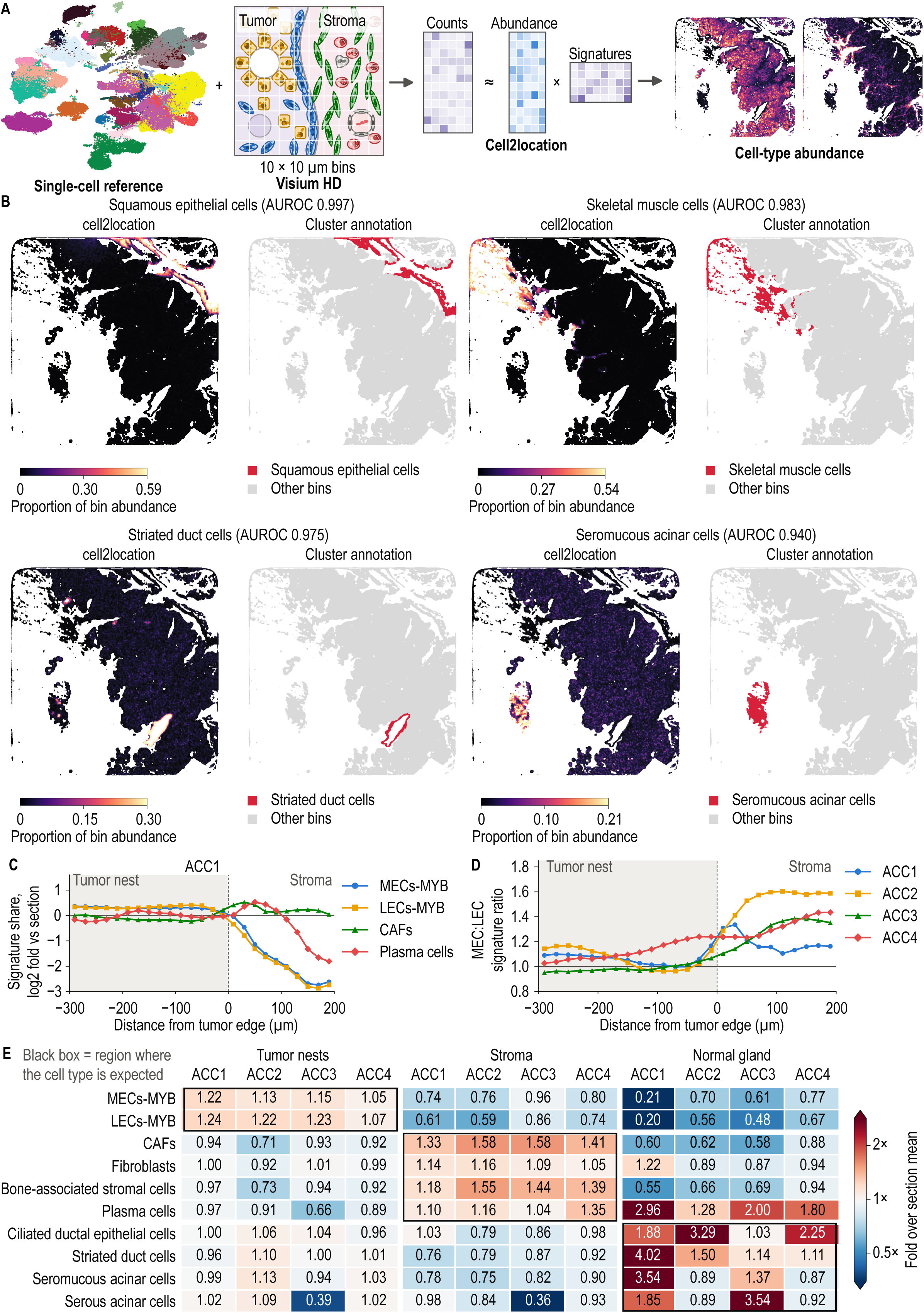
Cell2location maps malignant and non-malignant cell types in the SGACC tumor microenvironment. **(A)** Cell2location workflow. Cell-type signatures learned from the single-cell reference were used to estimate the abundance of each cell type in every 10 x 10 μm bin. **(B)** Cell2location estimates (left; proportion of the bin’s total abundance estimated by the model). Manual cluster annotation (right; red, bins annotated as that cell type) for squamous epithelial cells, skeletal muscle cells, striated duct cells and seromucous acinar cells in ACC1. AUROC, area under the receiver operating characteristic curve for separating the annotated bins from all other bins by the cell2location proportion. High AUROC values and matching location between manual and model localization highlights a well fitted model. **(C)** Cell2location share of myoepithelial cells-MYB (MECs-MYB), luminal epithelial cells-MYB (LECs-MYB), cancer-associated fibroblasts (CAFs) and plasma cells against signed distance from the tumor edge in ACC1 (negative inside the tumor nest), shown as log2 fold over the section mean in 20 µm distance bins. **(D)** Ratio of the MECs-MYB to the LECs-MYB share against distance from the tumor edge in ACC1 to ACC4. **(E)** Mean cell2location share of each cell type in the tumor nests, stroma and normal gland of each section, expressed as fold over the section mean. Black boxes mark the region in which each cell type is expected. Model predicts higher expression where it is predicted.

The reference-derived expression signatures closely reproduced the profiles of their respective cell types (Pearson r ∼ 0.80 across all cell types) **(Fig. S2B)**. The agreement of individual cell types ranged from r = 0.67 to 0.96 and was highest for the cell types that were most dominant in the tumor tissue sample and well represented in the single cell reference such as MECs-MYB; r = 0.96, CAFs; r = 0.94 and LECs-MYB; r = 0.93 **(Fig. S2D)**. To assess the quality of spatial mapping of cell types on tissue, the area under the receiver operating characteristic curve (AUROC) was calculated **(Fig. S2C)**. Interestingly the non-malignant compartments were localized with almost perfect accuracy. Squamous epithelial cells, skeletal muscle cells, striated duct cells and seromucous acinar cells were confined almost perfectly to their manually annotated regions with AUROC of 0.997, 0.983, 0.975 and 0.940 respectively **(Fig. 2B)**. Other normal glandular cell types were also resolved well, including serous acinar cells (AUROC = 0.95), and ciliated ductal epithelial cells (AUROC = 0.90 to 0.97). CAFs (AUROC = 0.83 to 0.93) and plasma cells (AUROC = 0.76 to 0.96) were also found to be consistently enriched in their annotated regions in spatial tumor samples. The spatial abundance maps of all the annotated cell types across all 4 samples are shown in (**Fig. S3).**

The two malignant cell types (MECs-MYB and LECs-MYB) were also enriched in their respective annotated regions although less sharply compared to the rest. This is expected as both the cell types have closely related expression patterns driven by MYB which is hard for the cell2location model to learn and differentiate perfectly. To orthogonally check if the localizations are reliable, we measured the mean proportions of these cell types as a function of distance from the tumor edge **(Fig. 2C, Fig. S4A)**. Both malignant signatures were highest in the tumor nests, ∼ 1.3-fold over the mean across all bins (section mean) and, these signatures dropped sharply across the boundary. Interestingly, in all the 4 SGACC tumor samples the MECs-MYB signature dropped slower than LECs-MYB with respect to distance **(Fig. 2D)**. The MEC:LEC ratio was 0.97 to 1.16 in the interior of tumor nest (60 to 160 μm from the edge). It rose to 1.08 to 1.24 at the edge (within 20 μm on either side) and reached 1.14 to 1.59 in the adjacent stroma (100 to 200 μm from the edge). This helps us to understand the tumor nest to be composed of more LECs-MYB in the center and MECs-MYB towards the periphery facing the stroma. Non-malignant cell types were depleted in the tumor nests. CAFs that are depleted in the tumor nest showed a sharp rise just outside the nests and this pattern was consistent across all SGACC tissues. This probably suggests that the tumor nests are surrounded by CAFs. The only immune cells that were identified at bin level are the plasma cells, showed an increase with increase in distance from the tumor edge. Except ACC1 that had peak plasma cells enrichment at ∼ 70 μm from the tumor edge, rest of the samples showed peak enrichment at 180 μm to 200 μm.

Finally, instead of looking at abundance at cell level we wanted to summarize the abundance maps at the level of tissue compartments. The tumor tissues were divided into tumor nests (MECs-MYB, LECs-MYB and stressed tumor cells – MYB), stroma (CAFs, plasma cells, stromal cells and bone-associated stromal cells) and normal glandular epithelium (normal gland; serous acinar, seromucous acinar, striated duct and ciliated ductal epithelial cells). The mean proportion of each cell type in each region was compared with section mean to estimate the enrichment in the expected region in the TME **(Fig. 2E)**. MECs-MYB and LECs-MYB were enriched in the tumor nest across all 4 SGACC tumor samples, giving us another level of validation that their spatial positioning in TME is accurate. The CAFs and the bone associated stromal cells were enriched in stroma and glandular cells like serous acinar and ductal epithelial cells were enriched in normal gland annotated tissue compartment corroborating the accuracy of cell2location constructed spatial maps of the SGACC TME.

Collectively, cell2location enabled cell type defined mapping of the single-cell reference onto the SGACC tumor sections at a resolution of 10 x 10 μm bins, providing estimates of both cell type abundance and spatial localization. As relatively few transcripts are captured per 10 x 10 μm bin, these estimates were interpreted as measures of relative spatial enrichment rather than absolute per-bin cell count. Within the limit of resolution, the single-cell reference constructed for the study reliably localized both the malignant and non-malignant components of SGACC TME and preserved the spatial relationship between the two malignant lineages and the surrounding stroma. These maps show a near cellular view of the SGACC TME.

### Spatially aware differential enrichment unravels pathways spatially restricted in SGACC TME across multiple tissue sections

Mapping the cellular composition of the SGACC tumor microenvironment allowed each tissue section to be partitioned into three compartments: tumor nest, stromal, and normal gland. To identify the gene expression programs specific to these compartment, raw counts from all bins within each compartment were summed per tissue section to generate a pseudobulk profile. PyDESeq2 (v0.5.4)[28,29] was implemented to find the differentially expressed genes between the compartments. The bins that are at the interface of tumor and surrounding stroma had expression signatures from both the regions as the bins might be composed of both cell types. To limit this mixing, we restricted the tumor vs stroma comparisons only to the core bins. These bins were located at least 80 μm away from the compartments being compared against **(Fig. 3A).** Of the 17,187 genes that were tested, 467 were significantly higher in the tumor core and 964 in the stromal core (FDR < 0.05, |shrunken log2FC| ≥ 0.5) **(Fig. S4B).** The stromal core had high expression of COL1A1, COL1A2, POSTN, SFRP2 hinting at a clear stromal program. The tumor core expressed MYB together with KIT and SOX4, two genes that are regulated by MYB and interestingly, two genes that are known to play important roles in ACC biology **(Fig. 3B, Fig. S4C)**. Next, we wanted to know the differentially expressed genes between the tumor nest and the normal epithelium. As the normal epithelium was spatially further away from the tumor nest there was no interfacial mixing hence, instead of limiting to the tumor core, the entire tumor nest was compared to the normal epithelium. Using the same significance threshold 919 genes were higher in tumor nest and 1,227 were higher in normal epithelium **(Fig. S4B)**. Like the tumor vs stroma comparison, MYB, SOX4, KIT, FABP7 and EN1 were significantly higher in the tumor nest **(Fig. S4D)**. The normal epithelium expressed secretory and ductal genes including BPIFB1, PIGR, SLPI and AQP3 **(Fig. 3C, Fig. S4D)**.

**Figure 3.**
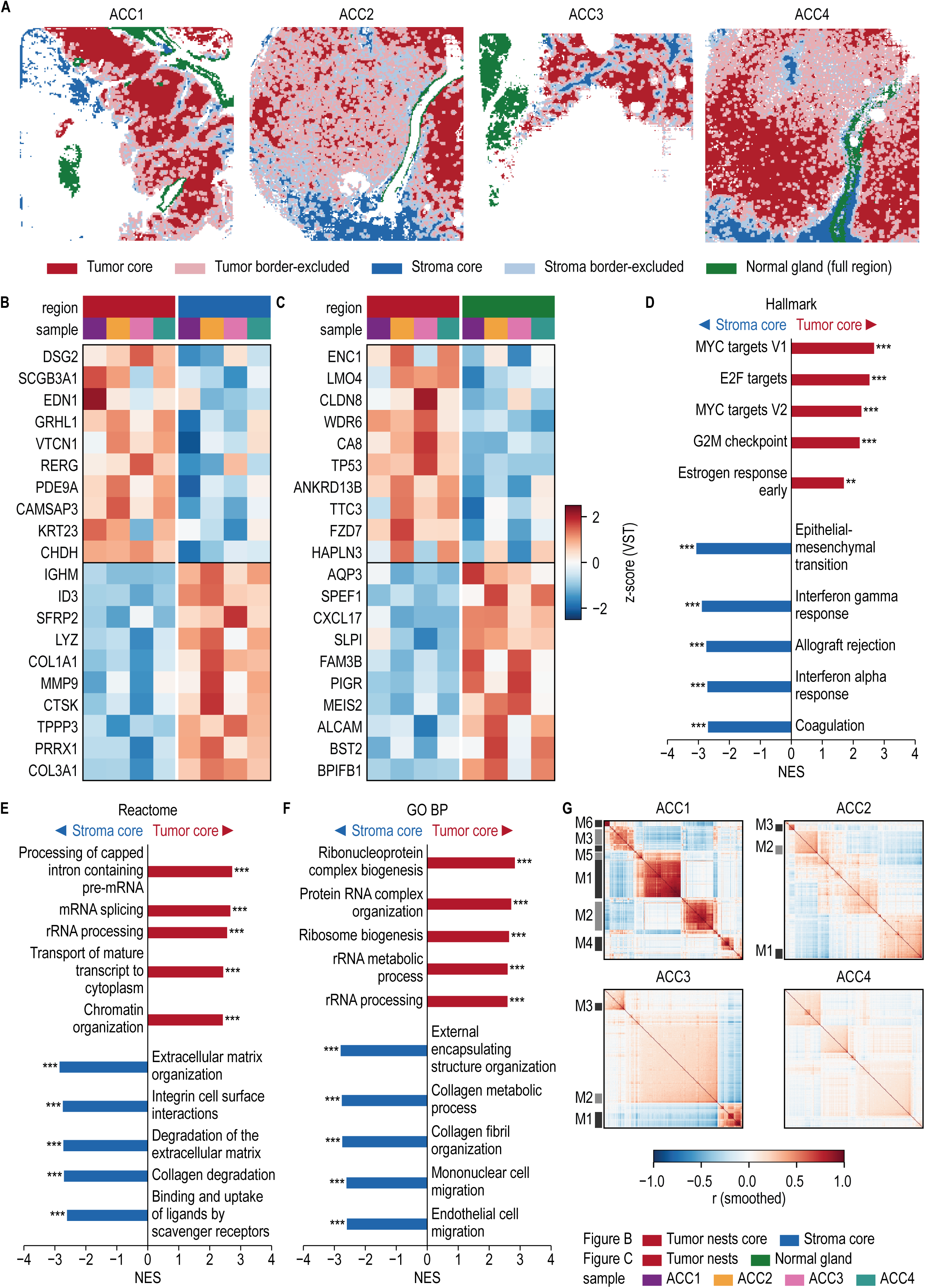
Pseudobulk differential expression and spatial autocorrelation separate tumor and stromal programs. **(A)** Tissue compartments used for pseudobulk differential expression on non-overlapping 50 µm tiles. Tumor core and stroma core tiles are defined as tiles that lie at least 80 µm from the other compartment. The border tiles were left out of the tumor versus stroma comparison. Normal gland used the full region as it didn’t border with tumor nests. **(B)** The 10 genes most significantly higher in the tumor core and the 10 most significantly higher in the stroma core (PyDESeq2, design ∼ section + region, n = 4 sections). Expression is variance-stabilized and z-scored per gene. **(C)** The same for tumor nests versus normal gland. **(D)** Hallmark gene sets enriched in the tumor core or the stroma core by pre-ranked gene set enrichment analysis (GSEA; genes ranked by the Wald statistic). The five most enriched biological programs with the largest absolute normalized enrichment score (NES) are shown for each side; a positive NES indicates enrichment in the tumor core. **(E)** Reactome and **(F)** GO Biological Process gene sets for the same comparison, shown as in (D). *** FDR q < 0.001; ** q < 0.01. **(G)** Smoothed gene-gene Pearson correlation of the 300 genes with the highest Moran’s I in each section, ordered by hierarchical clustering. Bars on the left mark the spatial gene modules (M1, M2, and so on). ACC4 formed no modules.

Next, we wanted to unravel potential pathways due to differential expression of these genes for which we performed gene set enrichment analysis using GSEAPy (v1.1.11)[30]. This showed that the tumor nests were enriched for proliferation associated programs such as Hallmark MYC targets V1, E2F targets, G2M checkpoint and ribosome biogenesis relative to both stroma and normal epithelium **(Fig. 3D-F; Fig. S4E-G)**. Interestingly the stroma was enriched for epithelial to mesenchymal transition (EMT) and extracellular matrix (ECM) organization terms **(Fig. 3D-F)**.

Normal epithelium was instead enriched for complement activation and cilium movement hinting at functional programs completely distinct from those of the tumor nest and stroma **(Fig. S4E-G)**.

To identify the spatially restricted gene expression patterns beyond the annotated compartments we computed Moran’s I spatial autocorrelation[31] for every gene in each tumor sample separately. To avoid gene expression, overlap due to small bins size of 10 μm, we made sure to use non-overlapping 50 μm bins. The number of spatially variable genes varied considerably across samples such as 475 in ACC1, 75 in ACC2, 71 in ACC3 and 22 in ACC4. COL1A1, COL1A2 and IGKC had high Moran’s I values in all the 4 SGACC TMEs, indicating spatially clustered, non-random expression of these genes. Next, we wanted to determine whether these spatially variable genes had groups of genes with similar expression patterns in its vicinity forming co-expression modules. We considered 300 genes ranked by Moran’s I in each sample and performed hierarchical clustering. This approach unraveled 7 modules in ACC1 and 3 each in ACC2 and ACC3 **(Fig. 3G, 4A; Fig. S5A-C)**. No modules were detected in ACC4 because of fewer SVGs due to lower sequencing depth of that section. Finally, to identify the modules conserved across tumor samples, the gene overlap between module pairs was tested. Out of 51 possible pairs only 7 pairs shared significantly more genes than expected by chance in the hypergeometric test (FDR < 0.05). These modules from different tumors based on presence of common genes were merged into groups that are conserved spatial programs across ACC tumor tissues in this study. We found 2 groups named G1 and G2 were, G1 comprised of ACC1 M3 and M7, ACC2 M2, ACC3 M2, whereas G2 comprised ACC1 M5, ACC2 M1 and ACC3 M3. G2 showed the strongest conservation as the individual modules shared 8 to 9 genes (FDR < 10^-18^) and COL1A1, COL1A2, COL3A1, FN1 and POSTN were hub genes in all three samples **(Fig. 4B, 4C)**. Apart from that interestingly, all three modules were enriched for same stromal and ECM related terms showing a conserved co-expression pattern across all tumor samples **(Fig. S5D)**. Comparatively modules in G1 overlapped less, sharing 2-4 genes per pair. G1 included a basal/myoepithelial program in ACC1, a luminal program in ACC2 and a module in ACC3 that is linked by shared ACC associated genes such as MYB and SOX4. Together these results show that the ACC tumor nests have spatially coherent MYB associated proliferative programs and the stroma has ECM programs. It also shows that few of the spatially coherent gene co-expression modules are conserved across multiple tumor tissues substantiating its relevance.

**Figure 4.**
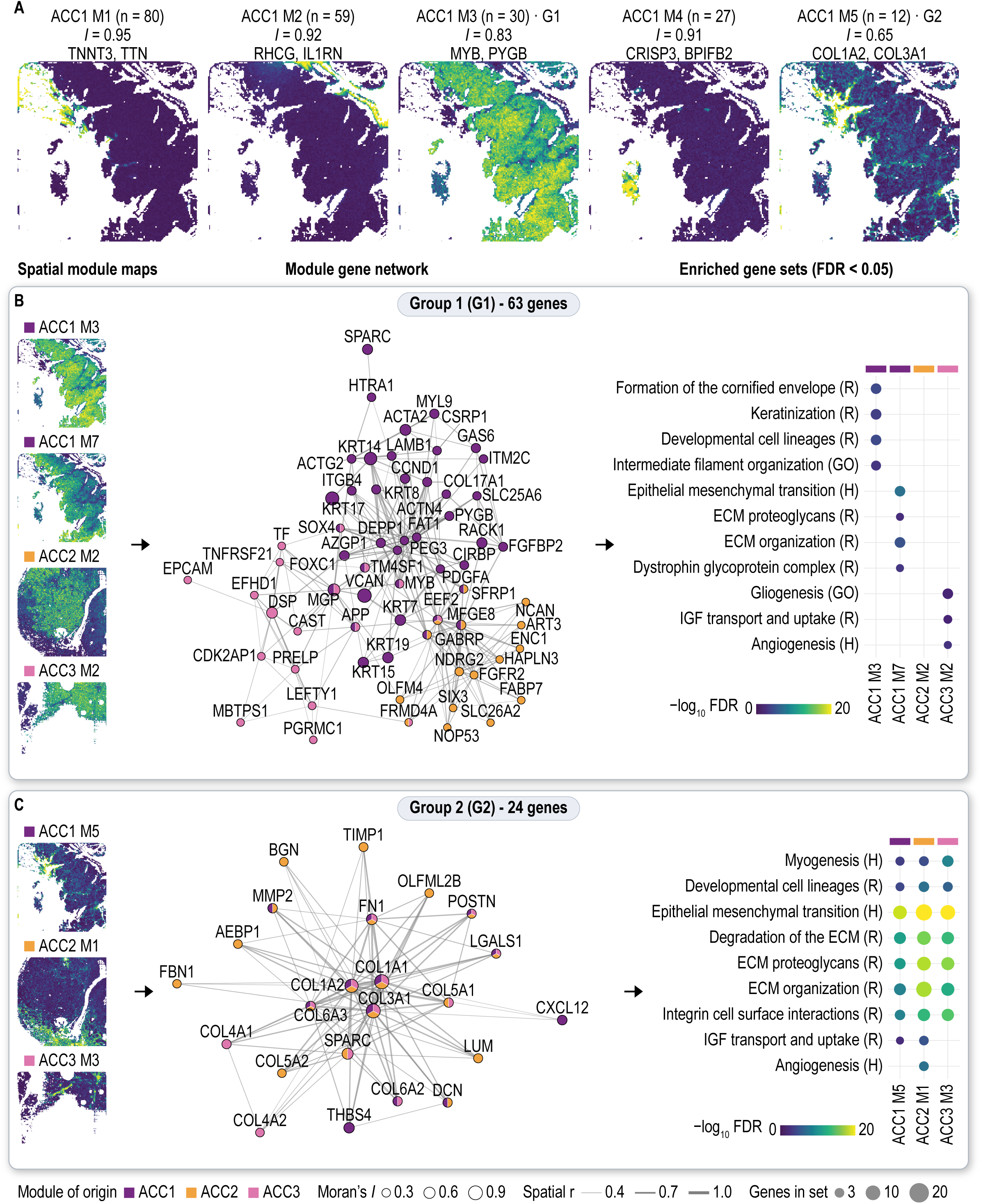
Spatial gene modules and the groups shared between tumors. **(A)** Score maps of the five largest spatial gene modules of ACC1 (mean z-score of the module genes in each 50 µm tile). Each title gives the number of genes (n), Moran’s I of the module score, the two hub genes and, where its located and the group its part of (G1 or G2). **(B)** Group G1 (63 genes), formed by modules from different sections with significant gene overlap (hypergeometric test, FDR < 0.05): score maps of its modules (left), co-expression network of their genes (center) and gene sets enriched in each module (right). Shows each modules being enriched for different terms. **(C)** Group G2 (24 genes), shown as in (B). In the networks, node color gives the module of origin, node size is proportional to Moran’s I, and edge width to the smoothed spatial correlation (r ≥ 0.4, up to five strongest partners per gene). In the dot plots, color gives −log10 FDR and dot size the number of module genes in the set. The three modules across different tissues enriched for the same pathway. H, Hallmark; R, Reactome; GO, GO Biological process.

### Single cell potency and spatial trajectory inference place MECs-MYB upstream of LECs-MYB and stressed tumor cells in SGACC

MECs-MYB and LECs-MYB were found to be the most abundant cell types in the SGACC TME in our 4-sample spatial dataset and in the reference single cell dataset made for this study. As these two cell types are transcriptionally very similar, we wanted to know if there is a shared transcriptional trajectory that can connect these two populations. We also asked how the cell types are positioned along this trajectory to determine which one arises first and how they are distributed across the SGACC TME. First, we implemented CytoTRACE2 (v1.1.0)[19] to determine the developmental potential of cell types in our single cell reference dataset. Then we rooted our Monocle3[32] trajectory analysis on the most potent cell type cluster to determine the trajectory. To avoid differences between studies and sequencing chemistries we used five ACC tumor samples from one whole-cell dataset (10x genomics 3’ sequenced, GSE342688) of our reference dataset. We choose the 5 samples based on the criteria that each MECs-MYB and LECs-MYB cells should make up more than 20% of the tumor cells, referred here as SC1-5 **(Table. S3)**. Depth matching and down sampling was done to make sure that the CytoTRACE2 scores are not influenced by the sequencing depth / number of detected genes. The samples we down sampled to 4,728 UMIs before scoring leaving 16,783 cells.

It was found that MECs-MYB had higher CytoTRACE2 potency scores than LECs-MYB in 4 / 5 single cell tumor samples **(Fig. 5A)**. The samples were pooled to test the significance using a linear mixed model that takes cell cycle scores as covariates and tumor as a random effect so that both baseline potency and the MEC-MYB / LEC-MYB difference can vary between the tumors **(Fig. 5B)**. This showed that the MECs-MYB scoring significantly higher than LECs-MYB (P-value = 0.011). The difference kept its direction and remained significant when cycling cell were removed (P = 0.012) and when each tumor leave one out analysis was performed (P ≤ 0.040). This helped us make the decision to use the MECs-MYB as the root for our trajectory analysis. Separate CytoTRACE2 runs for each tumor gave the same result, and depth matching removed the dependence of the score on transcript count **(Fig. S6B-D).** To trace the path between the two lineages, we applied Monocle3 after aligning the tumors and regressing out the cell-cycle scores. Out of the 8 clusters, two were made up almost entirely of MECs-MYB and expressed TP63, KRT5, KRT14 and ACTA2, while the luminal clusters expressed KIT, KRT7 and KRT8 **(Fig. 5C; Fig. S6E)**. The graph was rooted at the two MECs-MYB clusters, the pseudotime / trajectory ran through a mixed cluster (71% MECs-MYB) to a KIT⁺CD24⁺SOX4⁺ luminal cluster and then to mature luminal clusters **(Fig. 5C-G)**. Interestingly in 4 / 5 tumors the LECs-MYB potency dropped along pseudotime (Spearman ρ −0.40 to −0.54), which means the most distant LECs-MYB were least potent **(Fig. 5D-F).** Interestingly, the KIT⁺CD24⁺SOX4⁺ cluster had the highest mean potency of all clusters (0.48, against 0.44 and 0.41 in the two MECs-MYB clusters) and 78% of its cells are contributed by the sample SC5, the only one tumor sample that has LECs-MYB scored higher than MECs-MYB. Potentially, this cluster may be a luminal progenitor cell population with high expression of KIT and SOX4 that are two of the well-known MYB regulated genes highly expressed in tumor core **(Fig. S4C)**.

**Figure 5.**
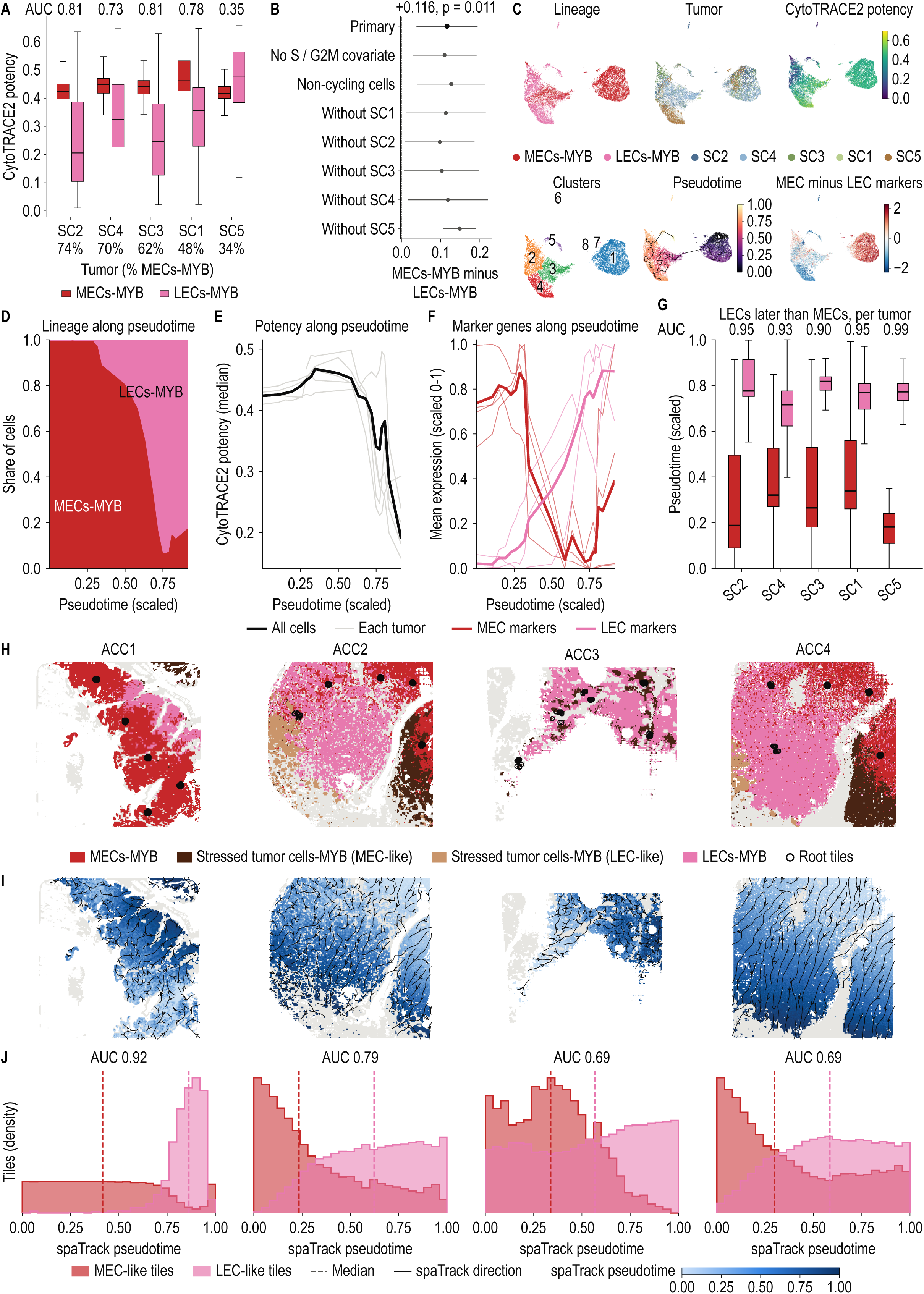
Single-cell potency and spatial trajectory inference place MECs-MYB upstream of LECs-MYB. **(A)** CytoTRACE2 potency of myoepithelial cells-MYB (MECs-MYB) and luminal epithelial cells-MYB (LECs-MYB) in five single-cell ACC tumors (SC1 to SC5; GSE342688) after down sampling to 4,728 UMIs per cell (16,783 cells, one pooled run; cells below this depth were removed). The percentage of MECs-MYB among tumor cells is given below each tumor and the AUC for MECs-MYB scoring higher than LECs-MYB above it. Boxes show the median and interquartile range. **(B)** Potency difference (MECs-MYB minus LECs-MYB) from a linear mixed model with S and G2M scores as covariates and tumor as a random effect (primary model, likelihood-ratio test), and from sensitivity models without the cell-cycle covariates, with non-cycling cells only, and with each tumor left out in turn. **(C)** UMAP of the pooled cells colored by lineage, tumor, CytoTRACE2 potency, Monocle3 cluster, Monocle3 pseudotime (black line, principal graph rooted in the MECs-MYB clusters) and MEC minus LEC marker expression (mean log expression of TP63, KRT5, KRT14 and ACTA2 minus that of KIT, KRT7 and KRT8). **(D)** Share of MECs-MYB and LECs-MYB along Monocle3 pseudotime. **(E)** Median CytoTRACE2 potency along pseudotime for all cells (black) and for each tumor (grey). **(F)** MEC markers (TP63, KRT5, KRT14, ACTA2) and LEC markers (KIT, KRT7, KRT8) along pseudotime, scaled 0 to 1; thin lines, single genes; thick lines, mean. **(G)** Monocle3 pseudotime of MECs-MYB and LECs-MYB in each tumor, with the AUC for LECs-MYB lying later than MECs-MYB. **(H)** Tumor nest tiles (50 µm) of ACC1 to ACC4 colored by group, with the 50 root tiles at five locations used for spaTrack. ACC3 has no MECs-MYB, so MEC-like stressed tumor cells-MYB served as the root there. **(I)** spaTrack pseudotime on the tissue; arrows show the direction of the ordering. **(J)** Distribution of spaTrack pseudotime in MEC-like tiles (MECs-MYB and MEC-like stressed tumor cells-MYB) and LEC-like tiles (LECs-MYB and LEC-like stressed tumor cells-MYB). Dashed lines show the medians and AUC shows the probability that a LEC-like tile lies later than a MEC-like tile.

As the robustness of the root and the root cell population is well inspected in the single cell dataset, we wanted to see the spread of these cellular states on the spatial tumor tissue for which we used spaTrack[33] implemented in python. We ran spaTrack on all 4 spatial tissues using the non-overlapping 50 μm bins from the Moran’s I analysis. Stressed tumor cells-MYB in each sample was sub clustered to find out their true identity marking them as MEC-like and LEC-like stressed tumor cells **(Fig. S6A)**. From the trajectory / pseudotime analysis of single cell reference dataset, the root was set as 50 MECs -MYB at 5 locations on the tissue in ACC1, 2, and 4. In ACC3 as there were no MECs-MYB, MEC-like stressed tumor bins were used **(Fig. 5H).** Interestingly the same direction was recouped in the spatial analysis as well with LEC-like bins being placed later in pseudotime compared to MEC-like bins (AUC 0.92, 0.79, 0.69 and 0.69 in ACC1 to ACC4) **(Fig. 5I, J)**. In ACC1, 2 and 4, LECs-MYB bins came after MECs-MYB bins and stressed tumor bins came last along the pseudotime **(Fig. S7A)**. The stressed fraction included the MEC-like stressed state that interestingly were placed after LECs-MYB despite the MEC-like expression profile along the pseudotime. Instead of bin level analysis we also checked the MEC / LEC marker gene levels along pseudotime. In ACC1, ACC2 and ACC4, the MEC markers TP63, KRT5, KRT14 and ACTA2 fell along pseudotime, whereas the LEC markers KIT, KRT7 and KRT8 rose or stayed level, as expected from the MEC-like root. As ACC3 didn’t have MECs-MYB and it was rooted at MEC-like stressed bins, they did not follow this pattern identically as rest of the samples **(Fig. S7B)**. Apart from just the marker genes we also wanted to check the programs enriched at the early intermediate and late states along the trajectory. The 3 samples with MECs-MYB as the start site put together showed enrichment for myoepithelial and basement membrane programs, including the Hallmark apical junction, epithelial–mesenchymal transition and myogenesis and Reactome extracellular matrix organization in the early third of the trajectory **(Fig. S7C, D).** The late third showed enrichment for mTORC1 signaling, unfolded protein response, and the E2F / G2M cell-cycle gene sets. Together, the potency and trajectory analysis gave us insights in both single cell and spatial datasets. Together this analysis showed that MECs-MYB have a higher potency than LECs-MYB in the reference single cell dataset that translated to capturing the same directionality on the spatial tissue along with determining the spatial spread of these cell types in the SGACC TME. The analysis also helped to capture the enrichment of specific programs along the trajectory helping us track the gene expression differences across the spatial tissue along a specific trajectory.

### SGACC tumors form structured millimeter scale territories of malignant cell types in TME instead of random cell type mixing

Trajectory analysis reveals the spatial distribution of the malignant cell types and how they connect to one another along the inferred trajectories. However, the trajectory is directed only by the neighboring bins with similar transcription profiles, so the analysis does not preserve the actual shape and scale of the malignant populations in TME. Hence, we wanted to know whether MECs-MYB, LECs-MYB and stressed tumor cells-MYB are intermixed with each other, or they segregate into structured territories and if so, how large are these territories. For this analysis we used the 50 μm bins and report distances of 100 μm or more. As a reference to compare against, random shuffling was done (1000 times) of the malignant cell type labels across the spatial tissue using SquidPy (v1.8.1)[34]. In the analysis it was found that each malignant cell population occupied large territories of the tissue which were absent from the shuffled spatial maps **(Fig. 6A; Fig. S8A)**. Across the four tumor samples, the largest territory was measured ranging 5.4 – 15.0 mm^2^ for MECs-MYB, 2.0 – 38.0 mm^2.^ for LECs-MYB and 0.66 - 9.4 mm^2^ for stressed cells. Interestingly, the largest territory in each of the 4 SGACC tumor samples were occupied by the dominant cell type (LECs-MYB in ACC2 to ACC4 and, MECs-MYB in ACC1). This shows that the dominant cell states in SGACC tumor which might be MECs or LECs are not randomly scattered instead, they for clear territories within the TME. In ACC4 the largest territory measured 7.5 mm² for MECs-MYB, 38.0 mm² for LECs-MYB and 9.4 mm² for stressed tumor cells-MYB. A 3 mm window enlarged the border between the stressed, LECs-MYB and MECs-MYB territories showing a sharp separation **(Fig. 6A, B)**. To actually measure how far the territories extend we calculated the percentage of tumor bins that are having the same state as the central bin at various distances from the center **(Fig. 6C)**. In ACC4, 61% of the bins 100 to 150 μm from a MECs-MYB bin were MECs-MYB, and 50% still were at 1 to 1.5 mm, against 23% by chance. Stressed tumor cells-MYB fell from 82% to 60% against 16% by chance, and LECs-MYB from 85% to 82% against 62% by chance. ACC2 and ACC1 behaved the same way, whereas the stressed bins of ACC3 fell to chance by about 1 mm (52% to 18% against 19%).

**Figure 6.**
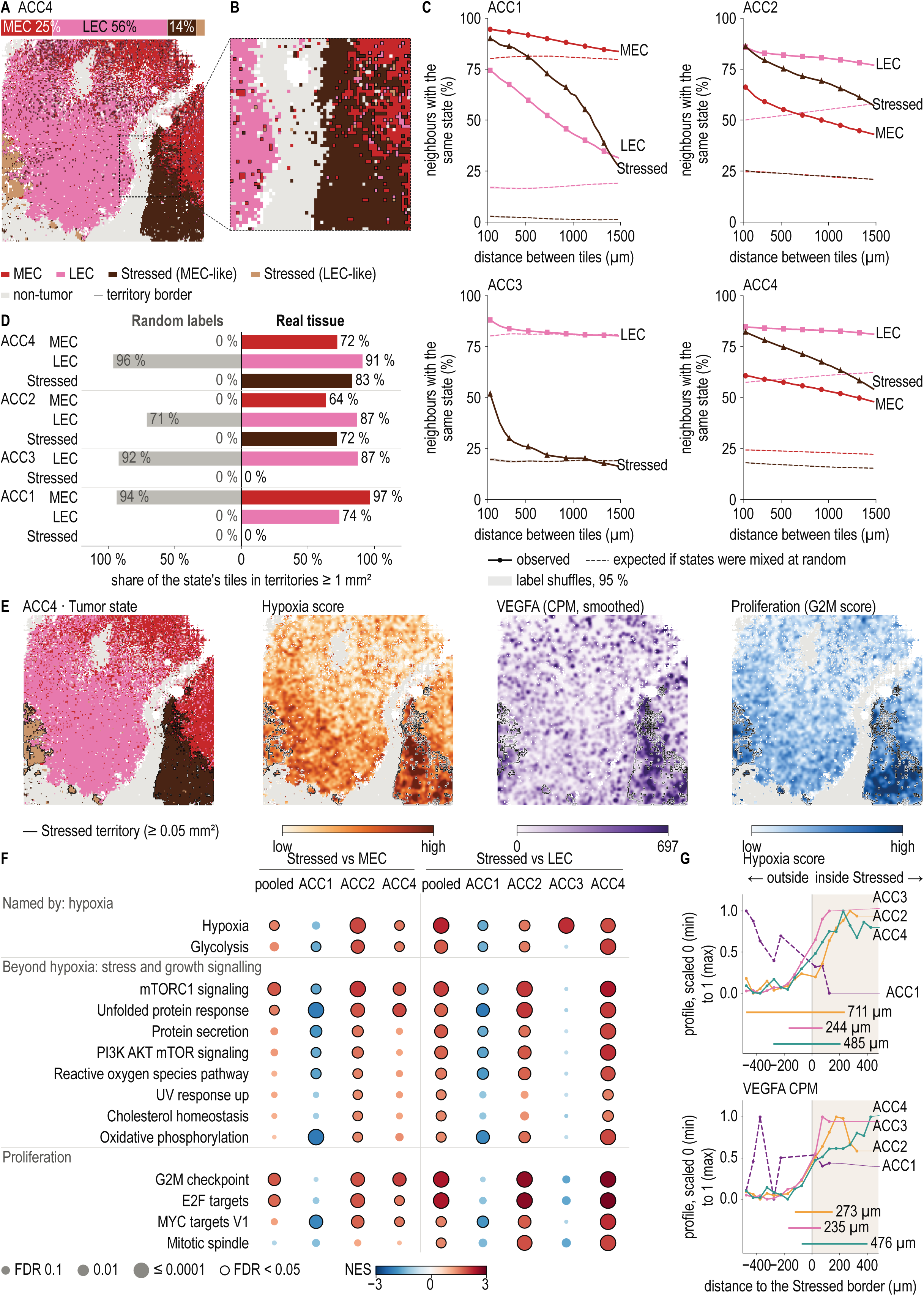
Tumor states form millimeter-scale territories, and the stressed territory carries hypoxia, growth and proliferation programs. **(A)** ACC4 tumor tiles (50 µm) colored by state, with territory borders. The bar above gives the share of each state among the tumor tiles, and the dashed box marks the 3 mm window enlarged in (B). **(B)** Enlarged ACC4 window with the borders between the LEC, stroma, stressed and MEC territories. **(C)** Percentage of tumor tiles at each distance that share the state of the center tile (solid lines), against the value expected under random mixing (dashed lines, the share of that state). **(D)** Percentage of each state’s tiles that lie in connected territories of at least 1 mm² in the tissue (right) and the upper 95% bound across 1,000 label shuffles (left, grey). States that make up more than about half of the tumor connect into one territory even when labels are random, so this panel is informative mainly for the minority states. **(E)** ACC4 maps of tumor state, hypoxia score, VEGFA expression in counts per million (CPM) and proliferation (G2M) score, smoothed with a Gaussian kernel (σ = 75 µm). Scores are univariate linear model scores of the Hallmark hypoxia and G2M checkpoint sets. Outlines mark stressed territories of at least 0.05 mm². **(F)** Hallmark GSEA of stressed tumor cells-MYB against MECs-MYB (pooled, ACC1, ACC2, ACC4) and against LECs-MYB (pooled, ACC1 to ACC4), based on PyDESeq2 pseudobulks of core tiles at least 100 µm from tiles of another label. Color gives NES, dot size FDR, and a black ring FDR < 0.05. **(G)** Hypoxia score and VEGFA CPM against signed distance to the stressed border (positive inside the territory), each section scaled from 0 (minimum) to 1 (maximum). Horizontal bars give the distance over which each profile rises from 10% to 90%; ACC1 (dashed) shows no rise. MEC, MECs-MYB; LEC, LECs-MYB; Stressed, stressed tumor cells-MYB, split into MEC-like and LEC-like subclusters.

To confirm these results, we also calculated co-occurrence ratio which hinted towards the same result **(Fig. S8B)**. In ACC1, ACC2 and ACC4 every state stayed above the 95% band that accounts for the random shuffles till 1.5 mm (permutation p-value = 0.001). A drastic difference was seen in stressed cell population in ACC3 in comparison to ACC1, 2 and 4 hinting that stressed cell territories were much smaller in ACC3 compared to the rest of the tumor samples. Apart from forming territories a cell population can look clustered only because it is filling a nest of its own. To check this the shuffling was repeated in each tumor nest. Almost all the cell populations stayed far above the null (permutation P = 0.001), except the stressed state of ACC1 as most of it was formed in a separate 0.9 mm^2^ section of the tissue in the corner. This analysis showed the presence of large cell state territories within the SGACC TME so, we wanted to know how much of each state lies in large territories. For this we joined the bins touching each other into patches[35] and for each malignant cell population we computed the percentage of its bins that fell in these patches of at least 1 mm^2^. It results hinted towards the same claim that the TME in SGACC is really structured **(Fig. 6D)**. Most of bins of malignant states were present in such patches such as, 72% of MECs-MYB bins and 83% of stressed bins in ACC4, 64% of MECs-MYB bins and 72% of stressed bins in ACC2 and 74% of LECs-MYB in ACC1. The stressed states of ACC1 and ACC3 did not form stressed patches of more than 1 mm^2^ hence were not included in calculation. In conclusion, all three states in ACC2 and ACC4, and MECs-MYB and LECs-MYB in ACC1, occupy territories larger than 1mm^2^ showing a well-structured arrangement of malignant states in SGACC TME.

Next, we wanted to explore what is happening in the stressed tumor cells-MYB as they are the 3^rd^ largest malignant cell population and an unanticipated cell population in the 4 SGACC tumor samples. These cells were named as they showed high expression of hypoxia related genes so, hypoxia and related response terms are expected to be high in that region. Hence, we wanted to explore what other biological processes are enriched in that region and how are they different from MECs-MYB and LECs-MYB population. Gene set enrichment analysis showed that, hypoxia, VEGFA and the G2M proliferation score were all high inside the stressed territories of ACC2 **(Fig. S8C)** and ACC4 **(Fig. 6E)**. In ACC3 the stressed patch was high for VEGFA and hypoxia **(Fig. S8D).** In ACC1 no enriched hypoxia or VEGFA programs were found, potentially because of its small size and low gene expression captured at the edge of the tissue **(Fig. S9A)**. It is also interesting to note that ACC1 is the only tumor tissue in the cohort that is MECs-MYB dominant and is the longest surviving patient **(Table S1)**. To test the enriched programs, we summed the counts of the core bins (at least 100 μm from bin of another type) per section and cell type. We then compared stressed tumor cells-MYB with MECs-MYB in ACC1, ACC2, and ACC4, and with LECs-MYB in all four sections, using PyDESEq2, with each section serving as a replicate. The significance threshold was kept at FDR < 0.05 and |apeGLM shrunk log2FC| ≥ 0.5 [36].

The stressed tumor cells-MYB against the LECs-MYB were enriched for the G2M checkpoint, E2F targets, hypoxia, EMT and mTORC1 and PI3K/AKT/mTOR signaling. Against MECs-MYB, the G2M checkpoint, mTORC1, E2F targets, the UPR and hypoxia were higher while EMT, myogenesis, complement and the interferon responses were lower **(Fig. 6F)**. Next, we checked in each section separately to see which sections carry these programs. In ACC2 and ACC4 mTORC1, UPR and G2M checkpoint were higher against both MECs-MYB and LECs-MYB. In ACC3 only hypoxia and EMT were higher, whereas in ACC1 these programs ran in the opposite direction. We also compared the LEC-like and MEC-like stressed cells with its LECs-MYB and MECs-MYB counterpart respectively **(Fig. S9B)**. Interestingly, related pathways were found to be enriched such as mTORC1, PI3K/AKT/mTOR, G2M and E2F in stressed LECs while stressed MECs showed significant enrichment of only mTORC1 and unfolded protein responses. This hints at the transcriptional similarity of the stressed cell types to their non-stressed counterparts. Stressed MECs might not be very different to MECs-MYB whereas stressed LECs show much more transcriptional variation when compared to LECs-MYB leading to more differentially enriched terms.

Finally, we wanted to know if the programs enriched in the stressed cells rises gradually from the boundary of the territory or is there an instant switch on / off, to estimate if the program is well contained within the stressed regions of the TME. To test this, we plotted the hypoxia score and VEGFA scores across the border and measured the distance over which each of them rose **(Fig. 6G)**. In ACC2, ACC3, and ACC4, both scores rose gradually over 244 to 711 μm for hypoxia and 235 to 476 μm for VEGFA. ACC1 interestingly showed no rise consistent with its lack of hypoxia and VEGFA enrichment. This showed that the cells towards the center of the territory are the most active, contributing to the enrichment of pathways that explains its functions. Altogether, these results showed that tumor states of SGACC are organized into huge millimeter-scale territories in three of the four sections instead of random mixing. ACC1 went the other way for most of the enriched programs maybe because it is the only tumor in this cohort that is not following a MYB-NFIB fusion but, one tumor sample is definitely not enough to link the two. Apart from that, ACC2 and ACC4 were two sections from the same tumor sample hence, both of them showing similar territory structures is not a surprise and is not considered as independent replicates. The millimeter scale territories shown in this study definitely warrants validation through spatially resolved analysis on a larger SGACC patient sample cohort to develop targeted therapies. To see whether these spatial patterns relate to clinical features, we compared the territory composition of each section with the clinical annotation **(Table S1)**. All four sections had a tubulo-cribriform pattern and stage III to IV disease, so histology and stage cannot explain the differences between them. The clearest differences were between patients with different outcomes. The only MEC-dominant section came from the longest-surviving patient (ACC1). In contrast, both sections from the patient who died (ACC2 and ACC4) were LEC-dominant and also contained the large proliferative, stressed territories as mentioned above. Finally, ACC3, came from another long-term survivor but, matched neither pattern: it had no MEC territory, only a small LEC territory and no large, stressed regions. As the comparison includes only three patients across four tissue sections, we could not test these patterns statistically and report them as descriptive observations.

### Malignant territories of SGACC show spatially restricted gene programs, while immune cells remain in stroma surrounding the tumor nest

The spatial co-occurrence analysis showed that MECs-MYB and LECs-MYB occupy millimeter-scale territories within the SGACC TME. Even though the territories are primarily composed of the same cell types we were skeptical if such large sections of tumor perform the same function. To test this, we did a Hotspot analysis[37] in python on the largest LECs-MYB and MECs-MYB territories of each of the 4 tumor samples **(Fig. 7A)**. Interestingly, this analysis showed the presence of several spatially restricted programs that were active at different regions within one territory. Across all 4 samples combined we had seven territories of 2.0 to 38.0 mm² (795 to 15,182 bins of 50 μm) as ACC3 only had LECs-MYB **(Fig. 7A)**. Our implementation of hotspot tested for autocorrelated genes in nearest bins (about 150 μm from each gene center) and groups the significantly autocorrelated genes into modules. We unraveled a total of 36 modules. As the bin size was set to 50 μm, it could potentially have 10 – 20 cells, so bins near boundary can have tissues can carry transcripts from adjacent territory. An elaborate multi step check was implemented to unravel the robust modules amongst them. A module made the cut only if its genes were not higher in a neighboring compartment (the other malignant cell type, stroma, plasma cells, muscle or normal gland), its score did not rise towards that compartment, it did not depend on sequencing depth, and it was not the hypoxia or immediate-early signal of a stressed border. After this only 9 of the 36 modules passed **(Fig. S11A).** Each unraveled module was named after the two most highly expressed genes. The ACC1 MECs-MYB territory showed four programs (MMP7/KRT5, GAS6/CALML5, KIT/BCAM and NFIB/NOTCH3), the ACC1 LECs-MYB territory had one (KRT7/AZGP1). LECs-MYB territories of ACC2 and ACC4 (two sections of same tumor) had two each while the MECs-MYB of ACC2 and ACC4 didn’t show any. Interestingly, no modules in LECs-MYB of ACC3 made the cut. Each of the highest scoring modules occupied 11 – 33% of bins in their respective territories across all tissues **(Fig. S10)**. Corroborating the credibility of the modules identified, ACC2 and ACC4 showed enrichment for the same programs (GABRP/MFGE8 and NCAN/VCAN) at similar locations with GABRP/MFGE8 programs sharing 10 genes and NCAN/VCAN sharing 7. The log2 enrichment of genes in respective programs is shown in **(Fig. 7B).** Altogether, this analysis showed interesting spatially restricted modules that are active at different regions within huge malignant territories in SGACC TME. No modules were conserved across tissue samples except the two between ACC2 and ACC4.

**Figure 7.**
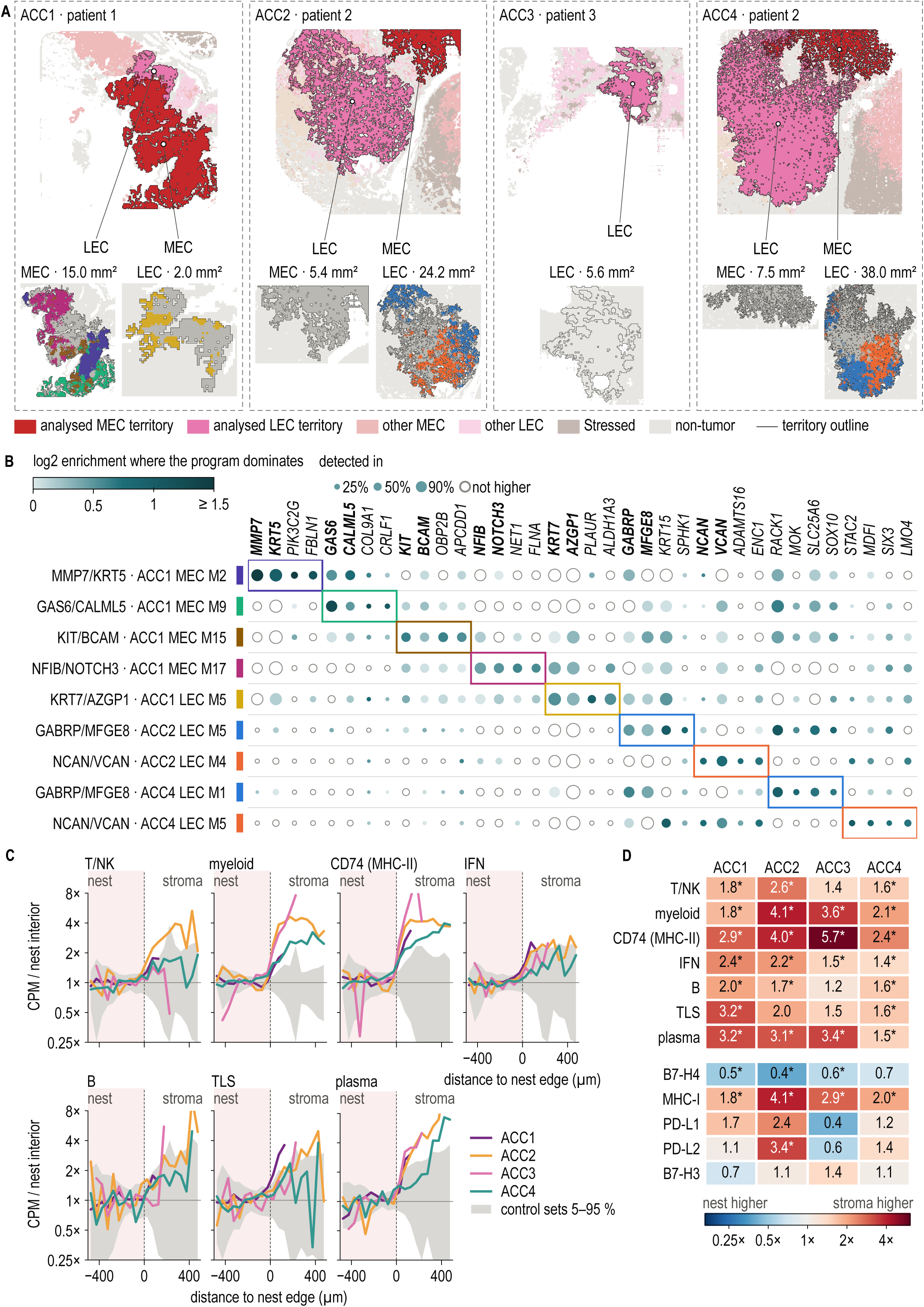
Spatial programs inside the tumor territories and immune exclusion at the nest edge. **(A)** The largest MEC and LEC territory of each section, filled and outlined on the whole tissue; other tumor tiles are faded. ACC2 and ACC4 are two sections of the same tumor (patient 2). Below each section, its territories are colored by the dominant credible Hotspot program (module scores z-scored within the territory; highest score when z ≥ 0.5). Grey tiles have no dominant program; ACC2 MEC, ACC3 LEC and ACC4 MEC had no credible program. **(B)** Genes of the nine credible Hotspot programs. Color gives the log2 enrichment of each gene in the tiles where the program dominates over the other tiles of the same territory (open circles, not higher), and dot size gives detection in the dominant tiles. Boxes mark each program’s own genes; bold genes name the program. **(C)** Immune gene-set expression signals against signed distance to the tumor nest edge (negative inside the nest, shaded), as pooled CPM in 50 µm distance bins relative to the nest interior (more than 100 µm from the edge). Grey band, 5th to 95th percentiles of 100 expression-matched control gene sets. CD74 alone represents MHC class II. **(D)** Exclusion index, the pooled CPM in stroma 50 to 300 µm from the edge divided by that in the nest interior, for immune and tumor-intrinsic gene sets. Asterisks, outside the 5th to 95th percentiles of the control sets. T/NK, T and natural killer cells; IFN, interferon response; TLS, tertiary lymphoid structure; B7-H4, VTCN1; MHC-I, B2M, TAP1, PSMB8 and PSMB9; PD-L1, CD274; PD-L2, PDCD1LG2; B7-H3, CD276; ACC1 to ACC4, patients.

The previous sections performed heuristic analyses on malignant nests, territories and also unraveled the programs enriched in them. The ACC tumors are frequently referred to as immune “cold”[38]. Hence, we wanted to localize the immune signals in the TME and see if any of them actually reach the malignant nests. Apart from plasma cells, other immune cells were too sparse to annotate at 10 x 10 μm bins **(Fig. 1D, E)** so, we decided to use the marker gene sets for T/NK cells, myeloid cells, B cells, tertiary lymphoid structures (TLS), plasma cells, interferon response (IFN) and CD74 (MHC-II), each compared with 100 expression-matched control sets. Across all 4 samples, marker gene sets for all 7 were higher in stroma than in tumor nests **(Fig. 7C)**. The ratio of expression in stroma 50 to 300 μm from the edge to expression in the nest interior was 1.4 to 2.6 for T/NK, 1.8 to 4.1 for myeloid, 2.4 to 5.7 for CD74 and 1.5 to 3.4 for plasma cells, and it exceeded the 95th percentile of the control sets in 24 of the 28 set and section combinations **(Fig. 7D)**. Altogether this shows that the immune transcriptional signal was therefore low overall. The signals were primarily concentrated in the stroma with a little signal in the tumor nest interior. Interestingly, this can be connected to the stromal enrichment of allograft rejection and interferon related terms shown in **Fig. 3D**. A low immune cell infiltration into the tumor nests can be attributed to lack of cellular machinery needed to present antigen to T cells. Hence, we wanted to check the levels of MHC class I antigen-presenting transcripts (B2M, TAP1, PSMB8 and PSMB9) across the tumor states. The transcripts were detected in MECs-MYB, LECs-MYB and stressed tumor cells in every section at similar levels to the normal gland in ACC1 and ACC3 and in a lower amount than normal in ACC2 and ACC4 **(Fig. S12A)**. Across the tumor-stroma boundary, MHC class I levels were considerably lower in the tumor nest than in the surrounding stroma **(Fig. S12C).** It should be noted that MHC class I was not completely depleted in the tumor nests. It can be assumed that the low T / NK signals within the nests could be because of the strong MHC signals located in the stroma overshadowing the nests.

Looking at this immune signal pattern, we wanted to further investigate if the tumor cells themselves are expressing any sort of inhibitory ligands that could limit immune infiltration. We profiled B7-H4 (VTCN1) across the tumor edge together with other B7 family ligands, such as PD-L1 (CD274), PD-L2 (PDCD1LG2) and B7-H3 (CD276). Interestingly, B7-H4 showed spatially opposite gene expression signals compared to immune signals **(Fig. S11B)**. It was highest in the nests and fell into the stroma with a ratio of 0.4 to 0.7 **(Fig. 7D, Fig. S12C).** In the nest interior B7-H4 was 1.3- to 5.4-fold higher than in normal gland in all four sections, the difference held after bins were down-sampled to equal depth, and it was highest in stressed tumor cells-MYB in ACC2, ACC3 and ACC4 **(Fig. S12A)**. In contrast, PD-L1 and PD-L2 were barely detected, and B7-H3 was as high in the stroma as in the nest, leaving B7-H4 as the only B7 ligand enriched in the tumor nest. Hence, B7-H4 can potentially be considered as a candidate regulator contributing to the confinement of immune signals to the stroma in these sections.

As plasma cells were the only immune population that was annotated in every tumor sample, we wanted to take a step further to see which antibody isotypes they produce. The isotype mix split by patient **(Fig. S12B)**. ACC1 (82% IgA, with more IgA2 than IgA1) and ACC3 (65% IgA) were dominated by IgA, whereas ACC2 and ACC4 were dominated by IgG1 (75%), and the two sections of this tumor agreed. As plasma cells were the predominant immune cells, we checked for immunoglobulin transcripts in both tumor nests and stroma. Immunoglobulin transcripts stayed at a low ambient level in tumor nests, indicating that plasma cells, even though in the majority, did not infiltrate into these nests. In conclusion, we saw that the SGACC tumor nests are indeed immune cold with the immune signals trapped in the stroma around it. To take a step further we also measured the immunoglobulin signals within the nests and stroma that lead to similar results. Even though the association was measured across 4 tissue sections, B7-H4 (VTCN1) signals showed inverse relation to immune signals across the tumor nests – stroma boundary potentially hinting at a real relationship which warrants validation in a larger cohort of this B7 family T cell inhibitory ligand.

## Discussion

In this study, we used Visium HD spatial transcriptomics together with an integrated multi-study single-cell reference to unravel a near-cellular map of the SGACC TME across four tissue sections from three patients. These maps showed that the two malignant lineages, MECs-MYB and LECs-MYB, follow a consistent arrangement at the edge of the tumor nest. It also showed that malignant cell states occupy millimeter-scale territories with their own different spatially restricted programs happening within them, and that immune signals remain in the stroma while B7-H4 (VTCN1) is enriched inside the tumor nests. The single-cell studies have already shown the pre-dominance of malignant myoepithelial and luminal epithelial populations in ACC[22,39] and our previous multi-omic study of 25 head and neck ACC tumors linked poor survival to the loss of myoepithelial tumor cells and the expansion of proliferative luminal states[40]. The four tumor sections from three patients analyzed in this study are independent of the previous cohort as they shared no patient, tissue section or sequencing runs between them. Spatial studies have also linked regional programs to pathological grade of the tumor and to stromal restriction of immune cells[11,41]. Our data implementing high resolution spatial transcriptomics add the scale and geometry of this organization together with spatially restricted gene expression programs, immune signal localization, and immunoglobulin isotope profiles across the SGACC TME.

Within the tumor nest we found that the MEC:LEC ratio increased from the tumor nest interior to the boundary. The CAF signatures rose sharply just outside the nests, placing MECs-MYB at the interface with a fibroblast rich stroma. This is consistent with the expected arrangement in which MECs-MYB sit on the basement membrane and LECs-MYB line the lumen. As both lineages are driven by MYB expression program, distinguishing between the two malignant cell types was difficult. The tumor core was also found to be expressing KIT and SOX4 along with MYB. This fits the model in which MYB drives distinct programs in each lineage, cooperating with TP63 in myoepithelial cells and with NOTCH signaling in luminal cells[42]. Previous studies have also shown that malignant myoepithelial cells express NOTCH ligands and luminal epithelial cells carry active NOTCH signaling[22]. The arrangement observed here, with LECs-MYB in the interior and MECs-MYB toward the exterior of the nests, provides a spatial setting in which this paracrine signaling could occur. Our potency and trajectory analyses placed MECs-MYB upstream of LECs-MYB. MECs-MYB scored higher than LECs-MYB with CytoTRACE2 in 4 of 5 tumors after depth matching and cell cycle adjustment in our single cell reference dataset, and the same direction was recovered on the tissue, where LEC-like bins lay later than MEC-like bins along spatial pseudotime in all four sections (AUC 0.69 to 0.92), and MEC markers declined while LEC markers rose or stayed level in the three sections rooted at MECs-MYB. This agrees with functional work in which myoepithelial-like ACC cells were more tumorigenic in xenografts and gave rise to ductal-like progeny through retinoic acid receptor signaling[43]. This pattern also agrees with our previous study that the tumor with good (> 5-year survival) and poor outcome (< 2-year survival) share a myoepithelial-like origin and differ in where they sit along the pseudotime trajectory with the poor outcome (more aggressive tumors) converging on a proliferative luminal state driven by MYC and E2F programs[40]. In our trajectory analysis on single cell reference data, it was observed that the KIT^+^CD24^+^SOX4^+^ luminal cluster had the highest mean potency of all clusters across all 5 tumors and its cells were contributed mostly (78%) from SC5, the one tumor in which LECs-MYB scored higher than MECs-MYB. This might be suggesting that the cluster may be a luminal progenitor-like state that is more prominent in some ACC tumors as well. Altogether, the potency and the trajectory analyses support a model in which the MECs-MYB occupy the upstream position, then LECs-MYB follow, and the proliferative stressed tumor cells-MYB form the end of the trajectory. In this trajectory, the late state was most extensive in the two sections from the diseased patients. Along with the MEC:LEC gradient from the nest edge to the interior, these results suggest that differentiation axes may be laid out in space across the tumor nest from the periphery to the core, and they provide a rationale for lineage tracing in patient-derived organoids or xenografts to test the direction of differentiation directly.

At a cluster level, MECs-MYB, LECs-MYB and stressed tumor cells-MYB did not mix randomly in the SGACC TME. Each malignant state was organized into territories several square millimeters in size. Territories of the same malignant state persisted over distances of up to 1.5 mm in 3 of the 4 tumor sections. These territories were absent from all 1,000 label-shuffled maps and, except for the ACC1 stressed state, persisted when labels were shuffled within each tumor nest. A multiregion sequencing study has shown branching evolution and regional genetic heterogeneity in ACC particularly in recurrent and metastatic disease[6] and a combined whole-exome and spatial profiling study has linked higher-grade regions to more proliferative expression states[41]. Even though the territories were made of same cell type populations, we found that the territories are not transcriptionally uniform. Our hotspot analysis unraveled programs that were active in different regions of the same territory. We found a total of 9 robust modules that were active in the malignant cell territories across the 4 samples. Modules such as NFIB/NOTCH3 and KIT/BCAM in the ACC1 MECs-MYB territory suggest that lineage and signaling programs vary within a single malignant state. The GABRP/MFGE8 and NCAN/VCAN programs recurred at similar locations in ACC2 and ACC4, but because these are sections of one tumor, this shows reproducibility within a tumor and not conservation across patients. The two sections also agreed in territory structure which supports the technical reproducibility of the measurements, but they are not treated as independent evidence. As both sections were cut from the same tumor, their similar territory makeup also suggests that these territories may extend through the depth of the tissue along the z-axis, although the distance between the two sections was not recorded and the depth they span therefore remains undetermined. No other active modules in the tumor nests were found to be shared between tumors. Our Moran’s I and spatial co-expression analysis revealed a stromal ECM module (outside tumor nests) built around COL1A1, COL1A2, COL3A1, FN1 and POSTN, conserved across 3 tumor sections from ACC1, ACC2 and ACC3. It was not detected in ACC4, potentially because of the lower sequencing depth of that section. This pattern hinted that the malignant programs in the tumor nests were largely tumor specific, but the stromal patterns were common. The stromal compartment was consistently found to be organized around the tumor nests. The stromal core showed high expression of COL1A1, COL1A2, POSTN and SFRP2, which leads to CAF related signatures just outside the nests. Previous studies have shown that ACC derived CAFs can lead the invasion of ACC cells by creating an invasive track through the matrix[44]. So, a matrix producing layer right around the tumor nest can potentially be linked to influence invasion, vascular supply and potential immune access. Further, the stressed territories added a layer of malignant cell type to the end of the inferred trajectory. In ACC2 and ACC4 that had huge stressed malignant cell type territories showed a combined hypoxia, VEGFA with mTORC1 and G2M/E2F programs. As these territories were huge, we measured the hypoxia and VEGFA scores and found they rose gradually towards the center from the territory border. In ACC, HIF-1α and other hypoxia-related proteins are increased and are prominent in necrotic areas and areas of perineural invasion[45] and hypoxia promotes VEGFA-dependent vasculogenic mimicry in experimental models[46]. The proliferative signal we found in hypoxic territories could be hinting that ACC cells do not respond to low oxygen in a single way, and that the balance between stress adaptation and proliferation may differ between regions and between tumor samples. These proliferative stressed territories may also relate to the discrete spatial niches of proliferative luminal cells that we previously observed in aggressive tumors[40]. The observed pattern was weaker in ACC3 and in ACC1 it went the opposite way. As the stressed population was annotated by taking into account the hypoxia-related genes, enrichment of hypoxia terms is partly expected hence, the enrichment of mTORC1, unfolded protein response and cell-cycle programs, together with the gradual rise from the border to the center, are the more informative findings in the stressed malignant territory.

Although the spatial sample set is small, the concentration of these programs in defined locations of the SGACC TME allows a few therapeutic hypotheses. The mTORC1 activity focused in the stressed territory is a candidate although everolimus produced disease stabilization without objective responses in a phase II ACC study[47]. High VEGFA in stressed territories are also relevant to VEGFR targeted treatment in such section of SGACC TME as lenvatinib and rivoceranib have shown activity in phase II trials and axitinib prolonged progression-free survival compared with observation[48–50]. With these high-resolution spatial samples we also looked into the spatial localization of the immune signals. All seven immune marker sets, including the T/NK, myeloid, CD74 and plasma cell signatures, were higher in the stroma than in the nests across the four sections, and immunoglobulin transcripts stayed at an ambient level in the tumor nests. The immune compartment in these tumors is present but held outside the malignant nests, showing a pattern of immune exclusion that is prevalent in SGACC TME making it immune cold. This pattern also agrees with the low numbers of tumor-infiltrating lymphocytes and the low or absent PD-L1 reported in ACC[10,51] and with the stroma restriction of immune cells by immunoprofiling[11]. Previous studies have shown that immunosuppressive macrophages dominate the immune compartment of salivary gland cancers[24] but, our marker-based signals did not resolve for such populations in the stroma or tumor nests of these samples. As plasma cells were the predominant immune population, we examined the isotypes they produced in each sample. IgA dominated in ACC1 and ACC3, and IgG1 in ACC2 and ACC4. In the immune excluded tumor nests, we saw that B7-H4 (VTCN1) stood out. VTCN1 was the only B7 family ligand that was enriched in the tumor nests as its spatial expression profile was exactly opposite to that of the immune signals. Interestingly, it was also noticed that VTCN1 was highest in the stressed malignant territory in 3 of the 4 samples. B7-H4 is an established feature of ACC: it is detected in most ACCs, and high expression is associated with solid histology and poor prognosis[52]. Spatial immunoprofiling linked B7-H4 to immune exclusion in aggressive ACC-I tumors[11], and single-cell profiling of salivary gland cancers localized VTCN1 to luminal-like ACC cells[24]. Our data placed VTCN1 in its tissue context, with an inverse relationship to immune signals across the boundary between nest and stroma and the highest expression in hypoxic, proliferative stressed territories.

This localization has direct therapeutic relevance. In a post hoc interim phase 1 analysis of 32 efficacy-evaluable patients with clinicopathologically defined aggressive ACC, emiltatug ledadotin (Emi-Le), an antibody-drug conjugate directed against B7-H4, produced an objective response rate of 46.9%, disease control in 81.3% and a median duration of response of 6.4 months[53]. Interestingly Emi-Le was administered on solid histology ACC tumors whereas all the tissues that are studied in this cohort are tubulo-cribriform but, they are still showing similar B7-H4 high tumor nests[52] hinting at a potential trial on other ACC tumor types as well. If B7-H4 protein follows the distribution we saw for VTCN1, the stressed territories may be especially exposed to them. Multiplex imaging of B7-H4 protein across luminal, myoepithelial and stressed territories would be a direct next step for developing a targeted analysis.

This study has limitations. The four samples came from three tumors as ACC2 and ACC4 are from same patient. Our section-level pseudobulk comparisons treated sections as replicates, so a larger cohort will be needed to substantiate these differences. . The 10 μm and 50 μm bins do not correspond to segmented single cells, and cell2location estimates depend on the composition of the reference and on capture efficiency[27], which is why we interpreted them as relative enrichment instead of absolute abundance of cell types. The two malignant lineages share MYB-driven expression patterns, which makes them harder to separate in the SGACC TME. The immune populations other than plasma cells were inferred from marker gene sets rather than annotated cells. The lower sequencing depth of ACC4 limited the detection of spatially variable genes and modules in that section.

## Conclusion

In conclusion, our high resolution spatial transcriptomic mapping of the SGACC TME helped us unravel the presence of several malignant territories of different scales ranging from μm² to mm². At a high resolution we could show that the LECs-MYB sit toward the interior and MECs-MYB toward a CAF-rich stroma, and the inferred trajectory runs from MEC-like toward LEC-like states. The huge malignant territories even though primarily made of one cell type show different spatially restricted programs within them. We observed a stressed territory that are hypoxic and proliferative that express higher levels of B7-H4 than other malignant territories. The immune transcriptional signals were concentrated in the stroma surrounding nests that expressed MHC class I antigen presentation transcripts, and signal levels were mapped across the boundary between stroma and tumor nests. This study, using the high-resolution spatial transcriptomic technology, shows the spatial lineage organization, millimeter-scale territories, hypoxic stress regions, and B7-H4-enriched, immune-excluded nests that provide a framework for validation in larger cohorts with protein-level and clinical data to determine which of these features drive recurrence and can be targeted therapeutically.

## Supplementary Information

Supplementary information document has all the supplementary tables and figure panels associated with this manuscript.

## Author contributions

G.B. [Conceptualization, Data curation, Formal analysis, Investigation, Methodology, Software, Validation, Visualization, Writing – original draft, Writing – review & editing]; K.H. [Investigation, Methodology, Writing – original draft, Writing – review and editing]; K.J.B. [Data curation, Validation, Writing – review and editing]; E.L.B. [Investigation, Writing – review & editing]; V.P. [Funding acquisition, Project administration, Resources, Supervision, Validation, Writing – original draft, Writing – review & editing]; J.S.E. [Conceptualization, Project administration, Investigation, Methodology, Resources, Supervision, Validation, Writing – original draft, Writing – review & editing].

## Institutional Review Board Statement

De-identified adenoid cystic carcinoma tumor samples were obtained from several institutions: the Department of Otorhinolaryngology and Maxillofacial Surgery, Zealand University Hospital; the Department of Otorhinolaryngology, Head and Neck Surgery and Audiology, Rigshospitalet; the Department of Pathology, Rigshospitalet, University of Copenhagen; and the Department of Ophthalmology, Rigshospitalet-Glostrup, University of Copenhagen, Copenhagen, Denmark. All samples were provided Formalin-Fixed and Paraffin-Embedded (FFPE) as 5-micron sections baked onto glass slides. Salivary gland samples with survival information had at least 5-year follow-up. All samples were collected in accordance with the principle of the Declaration of Helsinki and with Institutional Review Board-approved protocols: Danish Regional Ethics Committee (H-6-2014-086) and the Danish Data Protection Agency (Journal no. REG-94-2014).

## Informed Consent Statement

Informed consent was obtained from all subjects involved in the study.

## Data Availability Statement

The raw count matrices and annotated spatial data objects for all four SGACC tissue sections are publicly available in H5AD format together with the corresponding H&E-stained images with computational overlay in the Figshare repository at, **DOI: (The data generated in this study will be made publicly available through Figshare upon publication).**

Together with the analysis code, these files are sufficient to reproduce every analysis in this study from the primary – raw count data / matrices. Raw sequencing reads (FASTQ) are not available because they were lost in a cybersecurity incident affecting the storage servers. The single-cell reference used for cell type annotation was assembled from publicly available datasets deposited in the Gene Expression Omnibus under accession numbers GSE210171, GSE342688, GSE217084 and GSE294017, and from the Tabula Sapiens – Salivary Gland [The Tabula Sapiens Consortium et al. (2024) bioRxiv].

All codes / scripts used for data processing and analysis along with an interactive website enabling the visualization of spatial plots directly on the tissue is available on GitHub (https://github.com/Gopikrishnan-Bijukumar/SGACC-VisiumHD-spatial-analysis).

## Acknowledgements

The authors thank the Purdue University Rosen Center for Advanced Computing for providing Anvil computational resources funded by the National Science Foundation (Grant Number: 2005632). Access to computing resources was provided through the Advanced Cyberinfrastructure Coordination Ecosystem: Services & Support (ACCESS) program.

Anthropic Claude was used for language refinement and grammatical editing. Anthropic Claude was used to assist in generating analysis code in Python and R. All generated code was manually reviewed and validated by the authors. The authors reviewed, edited, and verified all AI-assisted text and code and take full responsibility for the content of the manuscript.

