## Supplementary Information (Tables S1-S3, Figures S1-S12) for "High-Resolution Spatial Transcriptomics Reveals Millimeter-Scale Malignant Territories and Immune Exclusion patterns in Human Salivary Gland Adenoid Cystic Carcinoma"

**Supplementary Table 1 (Table S1):** Clinical and molecular characteristics of salivary gland adenoid cystic carcinoma (SGACC) samples analyzed using high resolution spatial transcriptomics chips.

| Sample | Mapping | Translocation | Vital Status | Survival [Month] | Gender | Histological Form | Stage |
|---|---|---|---|---|---|---|---|
| STST02 | ACC1 | MYB8-Other | Alive | >204 | Male | Tubulocribriform | III–IV |
| STST03 | ACC2 | MYB12-NFIB | Deceased | <125 | Female | Tubulocribriform | III–IV |
| STST05c37 | ACC3 | MYB14-NFIB | Alive | >175 | Male | Tubulocribriform | III–IV |
| STST08-c49 | ACC4 | MYB12-NFIB | Deceased | <125 | Female | Tubulocribriform | III–IV |

**Supplementary Table 2 (Table S2):** Publicly available single-cell and single-nucleus RNA sequencing datasets of adenoid cystic carcinoma and healthy salivary gland used as reference samples.

| Donor | Tissue Site | Histology | Technology | Samples | Cells | Malignant Cells |
|---|---|---|---|---|---|---|
| **Parikh et al., 2022 (Cell Rep)** *GEO GSE210171 · ACC* | | | | | | |
| ACC19 | Left parotid gland | Tubular/solid | Smart-seq2 | 3 | 124 | 93 |
| ACC21 | Right base of tongue (minor salivary) | Cribriform | Smart-seq2 | 3 | 163 | 118 |
| ACC22 | Left floor of mouth (minor salivary) | Cribriform | Smart-seq2 | 3 | 200 | 30 |
| ACC2 | Right preauricular (parotid region) | Tubular/cribriform | Smart-seq2 | 3 | 262 | 93 |

| Donor | Tissue Site | Histology | Technology | Samples | Cells | Malignant Cells |
|---|---|---|---|---|---|---|
| ACC5[a] | Left parotid gland | Cribriform | Smart-seq2 | 4 | 309 | 257 |
| ACC7 | Right buccal mucosa (minor salivary) | Tubular/solid | Smart-seq2 | 4 | 314 | 106 |
| **Zhou et al., 2023 (Exp Mol Med)** *GEO GSE217084 · ACC* | | | | | | |
| Patient A[b] | Salivary gland (site not specified) | – | 10x 3′ | 2 | 16,241 | 3,957 |
| **Zuljan et al., 2025 (Nat Commun)** *GEO GSE294017 · ACC* | | | | | | |
| P32 | Salivary gland (site not specified) | – | 10x 3′ (single nucleus) | 1 | 4,593 | 4,460 |
| **Yuan et al., 2026 (preprint)** *GEO GSE342688 · ACC* | | | | | | |
| ACC-1 | Sinonasal/skull base (minor salivary glands) | – | 10x 3′ | 1 | 8,944 | 6 |
| ACC-2 | Sinonasal/skull base (minor salivary glands) | – | 10x 3′ | 1 | 6,966 | 5 |
| ACC-3 (SC1)[c] | Sinonasal/skull base (minor salivary glands) | – | 10x 3′ | 1 | 6,030 | 922 |
| ACC-4 | Sinonasal/skull base (minor salivary glands) | – | 10x 3′ | 1 | 7,202 | 4,195 |
| ACC-5 (SC2)[c] | Sinonasal/skull base (minor salivary glands) | – | 10x 3′ | 1 | 8,059 | 3,874 |
| ACC-6 (SC3)[c] | Sinonasal/skull base (minor salivary glands) | – | 10x 3′ | 1 | 7,629 | 3,211 |
| ACC-7 (SC4)[c] | Sinonasal/skull base (minor salivary glands) | – | 10x 3′ | 1 | 6,558 | 6,325 |
| ACC-8 (SC5)[c] | Sinonasal/skull base (minor salivary glands) | – | 10x 3′ | 1 | 7,700 | 4,357 |
| **Tabula Sapiens Consortium, 2022 (Science)** *CELLxGENE · Normal* | | | | | | |
| TSP14 | Parotid gland; sublingual gland | – | 10x 3′; Smart-seq2 | 2 | 17,607 | 0 |
| TSP21 | Parotid gland; submandibular gland | – | 10x 3′ | 1 | 3,755 | 0 |

| Donor | Tissue Site | Histology | Technology | Samples | Cells | Malignant Cells |
|---|---|---|---|---|---|---|
| TSP25 | Sublingual gland; submandibular gland | – | 10x 3′ | 1 | 7,465 | 0 |
| TSP7 | Parotid gland | – | 10x 3′; Smart-seq2 | 2 | 9,068 | 0 |
| **Total: 5 datasets, 20 donors (16 ACC, 4 healthy)** | | | | **37** | **119,189** | **32,009** |

**Supplementary Table 3 (Table S3):** Publicly available single-cell RNA sequencing adenoid cystic carcinoma tumors (SC1–SC5) included in this study and their malignant cell composition.

| Tumor ID | Source Sample ID | Malignant Cells (n) | MECs-MYB (n) | LECs-MYB (n) | MECs-MYB Share (%) |
|---|---|---|---|---|---|
| SC1 | ACC-3 | 922 | 463 | 459 | 50.2 |
| SC2 | ACC-5 | 3,874 | 2,865 | 1,009 | 74.0 |
| SC3 | ACC-6 | 3,211 | 2,088 | 1,123 | 65.0 |
| SC4 | ACC-7 | 6,325 | 4,541 | 1,784 | 71.8 |
| SC5 | ACC-8 | 4,357 | 1,766 | 2,591 | 40.5 |
| **Total** | **–** | **18,689** | **11,723** | **6,966** | **62.7** |

Supplementary Figure Panel 1

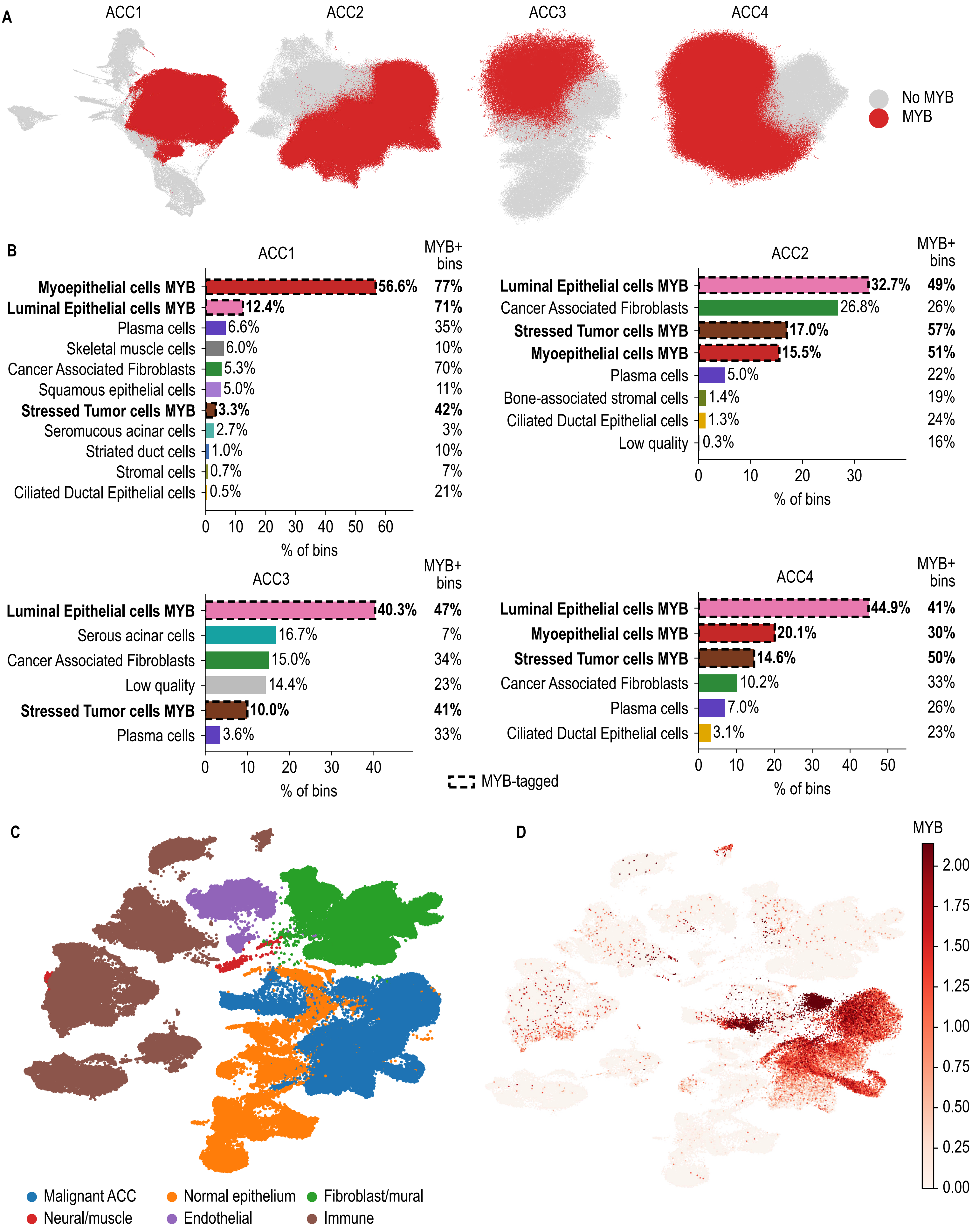

**Figure S1. MYB tagged regions, cell-type composition of each section and compartments of the single-cell reference. (A)** UMAP of each section with bins of MYB-tagged clusters in red. An epithelial cluster was tagged when its MYB detection through the heuristic approach when, compared with about 200 expression-matched control genes as an odds ratio, exceeded the spillover level of the non-epithelial bins of that section and persisted in bins away from other tumor clusters. **(B)** Cell-type composition of each section as the percentage of bins. Dashed outlines and bold labels mark MYB-tagged cell types; the right column gives the percentage of bins in each cell type with at least one MYB count. **(C)** UMAP of the single-cell reference colored by compartment. **(D)** MYB expression (log1p counts per 10,000) on the reference UMAP.

### Supplementary Figure Panel 2

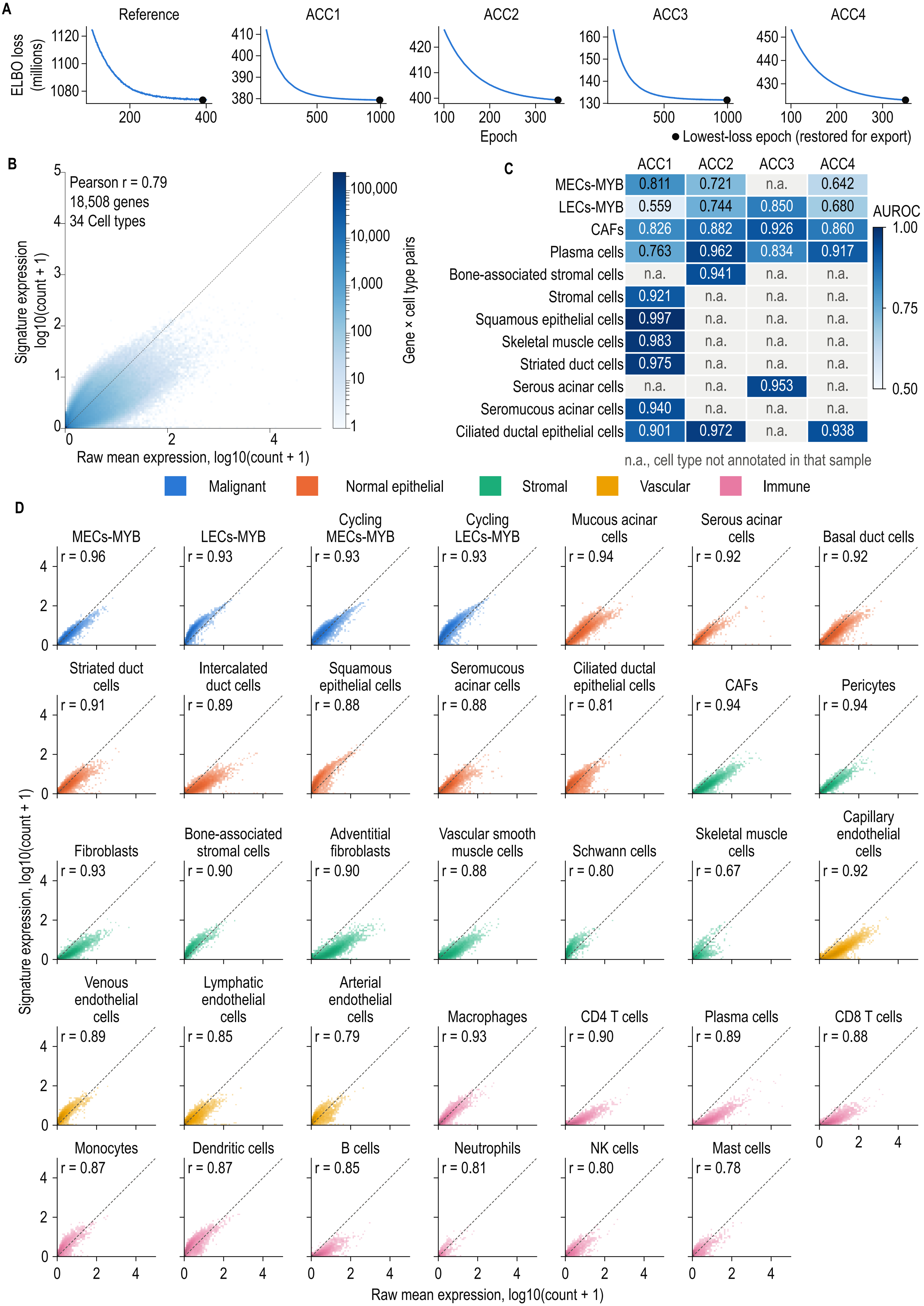

**Figure S2. Quality control of the cell2location model. (A)** Training loss (evidence lower bound, ELBO) along epochs for the reference regression model and the spatial mapping models of ACC1 to ACC4; the first 100 epochs are not shown. Black dots mark the lowest-loss epoch, which was restored for export. Shows characteristic curve signatures without much noise, hinting at clean training steps. **(B)** Reference signatures against the raw mean expression of the same cells for every gene and cell-type pair (18,508 genes, 34 cell types), on a log10(count + 1) scale. The color aligned with the dotted line shows accordance and not a lot of deviation. **(C)** AUROC of each annotated cell type in each section, computed from the cell2location proportion against the cluster annotation. n.a., cell type not annotated in that section. **(D)** As shown in (B), for each reference cell type separately, colored by compartment, with the Pearson r on the log scale.

Supplementary Figure Panel 3

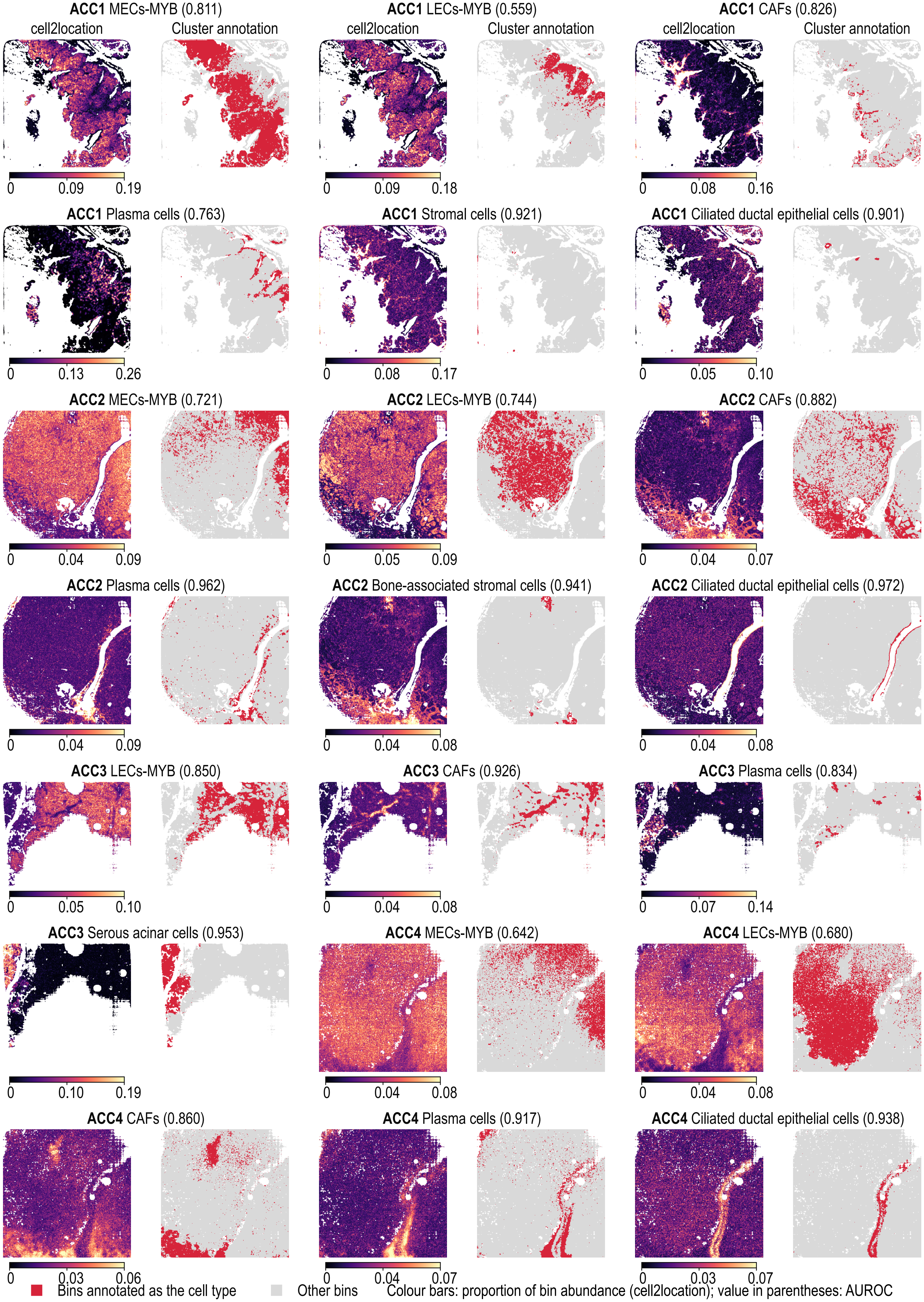

**Figure S3. Cell2location maps against the cluster annotation for the remaining cell types.** For each section (ACC1 to ACC4), the cell2location proportion (left of each pair; proportion of the bin's total abundance estimated by Cell2Location model) is shown beside the bins manually annotated as that cell type (right; red). Values in parentheses are AUROC. MECs-MYB and LECs-MYB include their cycling states. The four ACC1 cell types of Figures 2B are not shown here.

Supplementary Figure Panel 4

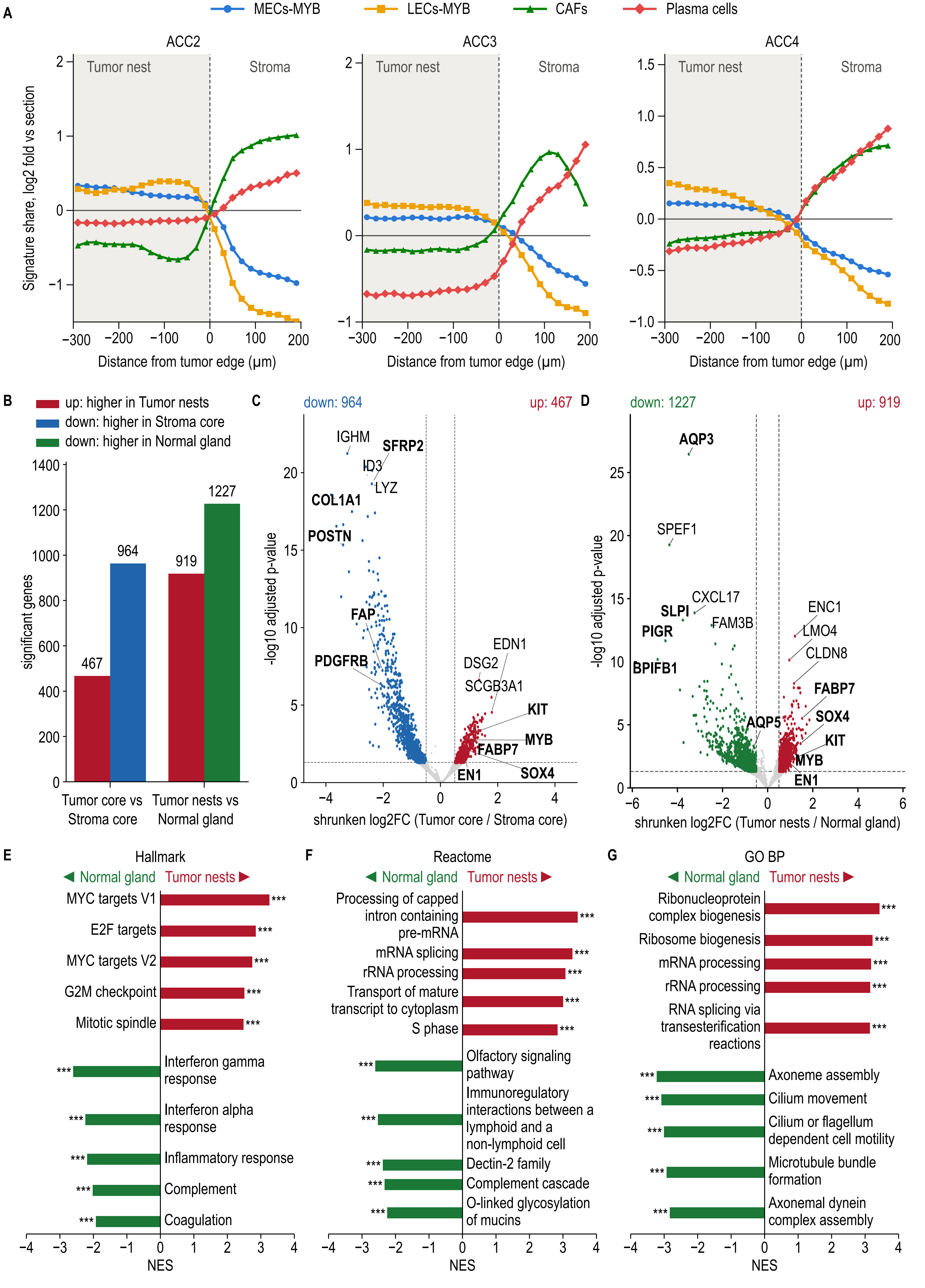

**Figure S4. Cell2location boundary profiles and differential expression between tumor nests, stroma and normal gland. (A)** Cell2location share of MECs-MYB, LECs-MYB, CAFs and plasma cells against signed distance from the tumor edge in ACC2, ACC3 and ACC4, shown as in Figure 2C for ACC1 **(B)** Number of differentially expressed genes (FDR < 0.05 and absolute apeGLM-shrunken log2 fold change ≥ 0.5) for tumor core versus stroma core and for tumor nests versus normal gland. **(C)** Volcano plot of tumor core versus stroma core. Dashed lines mark the significance thresholds. Bold labels are curated (well-known) ACC and stromal marker genes; the other labels are the top genes by adjusted P value. **(D)** Volcano plot of tumor nests versus normal gland, shown as in (C). **(E)** Hallmark, **(F)** Reactome and **(G)** GO Biological Process gene sets enriched in tumor nests or normal gland (GSEA), shown as in Figure 3D. *** FDR q < 0.001.

**Supplementary Figure Panel 5**

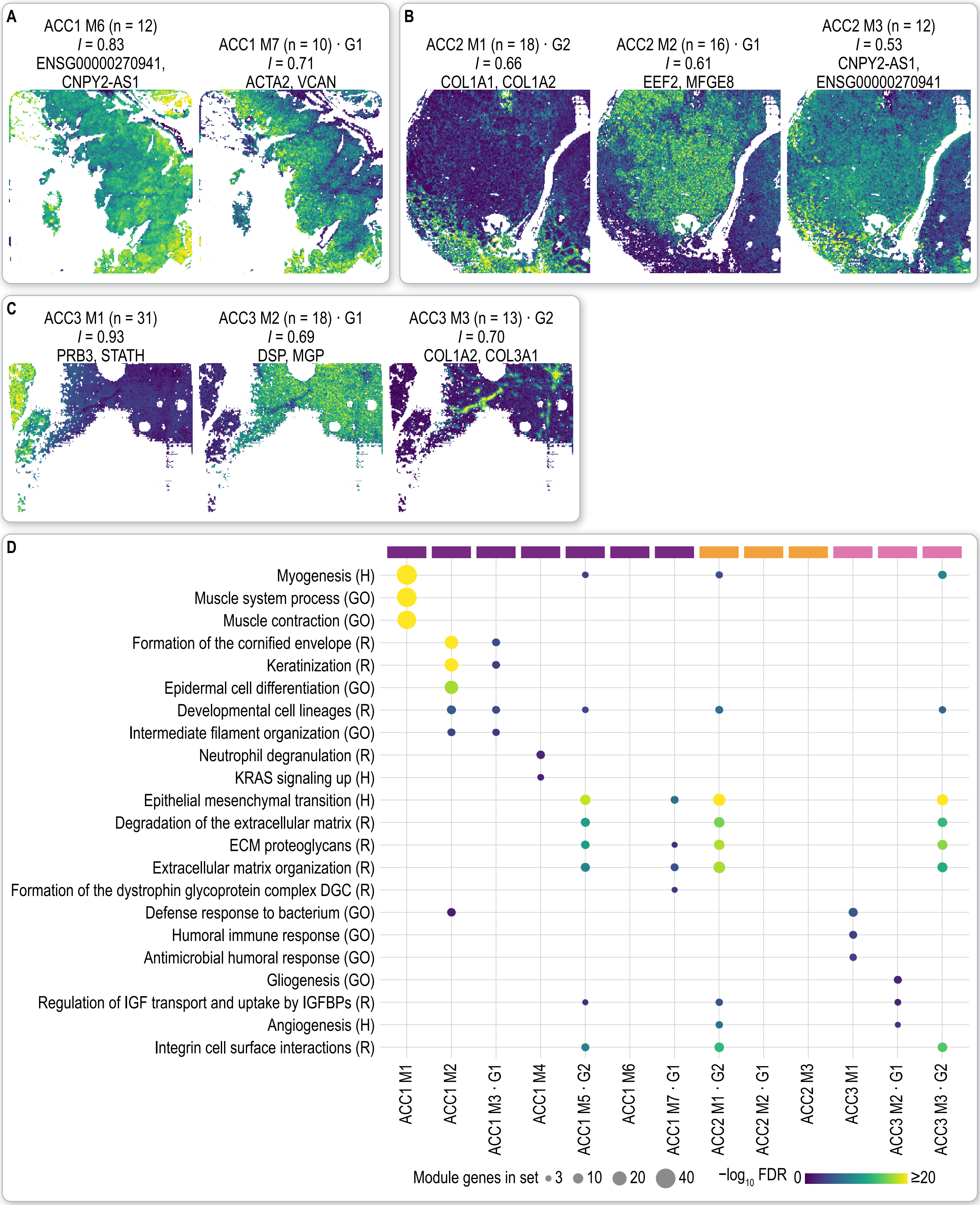

**Figure S5. The remaining spatial gene modules and their enriched gene sets. (A)** Score maps of ACC1 modules M6 and M7. **(B)** Score maps of ACC2 modules M1 to M3. **(C)** Score maps of ACC3 modules M1 to M3. Maps and titles are shown as in Figure 4A. ACC1 M6 and ACC2 M3 share a smooth gradient of mostly unannotated transcripts across the capture area and were regarded as technical. **(D)** Gene sets enriched in each module (hypergeometric test; FDR < 0.05 with at least three module genes). Color gives −log10 FDR (capped at 20) and dot size the number of module genes in the set. G1 and G2 mark the modules of the matched groups. H, Hallmark; R, Reactome; GO, GO Biological Process.

Supplementary Figure Panel 6

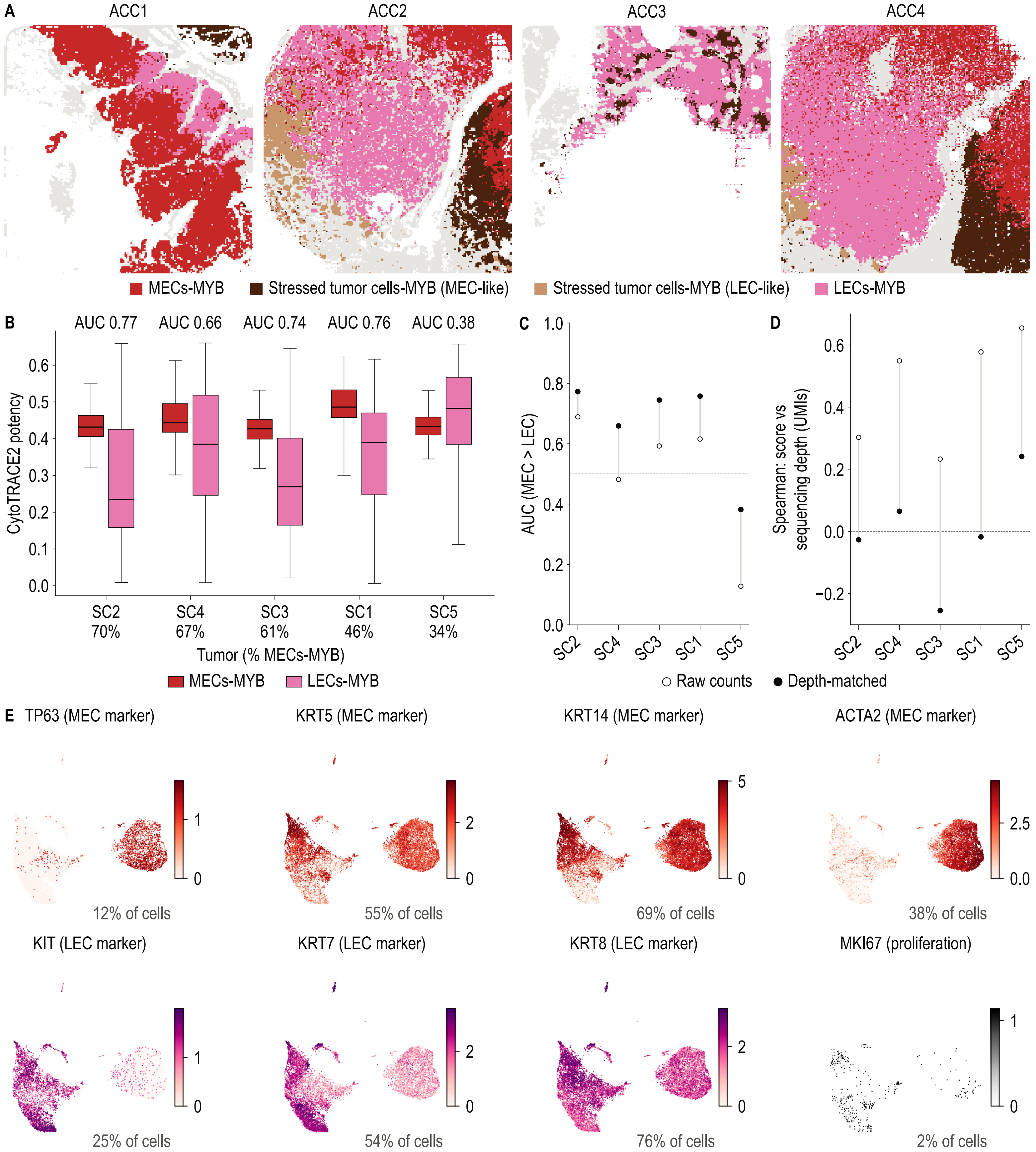

**Figure S6. Tumor groups for spaTrack, per-tumor CytoTRACE2 runs and lineage markers. (A)** Tumor nest tiles of ACC1 to ACC4 colored by MECs-MYB, LECs-MYB and the MEC-like and LEC-like subclusters of stressed tumor cells-MYB. **(B)** CytoTRACE2 potency of MECs-MYB and LECs-MYB from a separate run with depth matched cells for each single-cell tumor (SC1 to SC5), with the AUC for MECs-MYB higher than LECs-MYB above and the percentage of MECs-MYB below. Boxes as in Figure 5A. **(C)** AUC of each tumor from raw counts (open circles) and from depth-matched counts (filled circles). Dotted line represents AUC = 0.5. **(D)** Spearman correlation between potency score and sequencing depth (UMIs) in each tumor, from raw and depth-matched counts. **(E)** MEC markers (TP63, KRT5, KRT14, ACTA2), LEC markers (KIT, KRT7, KRT8) and MKI67 on the UMAP of Figure 5C. Percentages give the share of cells in which the gene was detected.

Supplementary Figure Panel 7

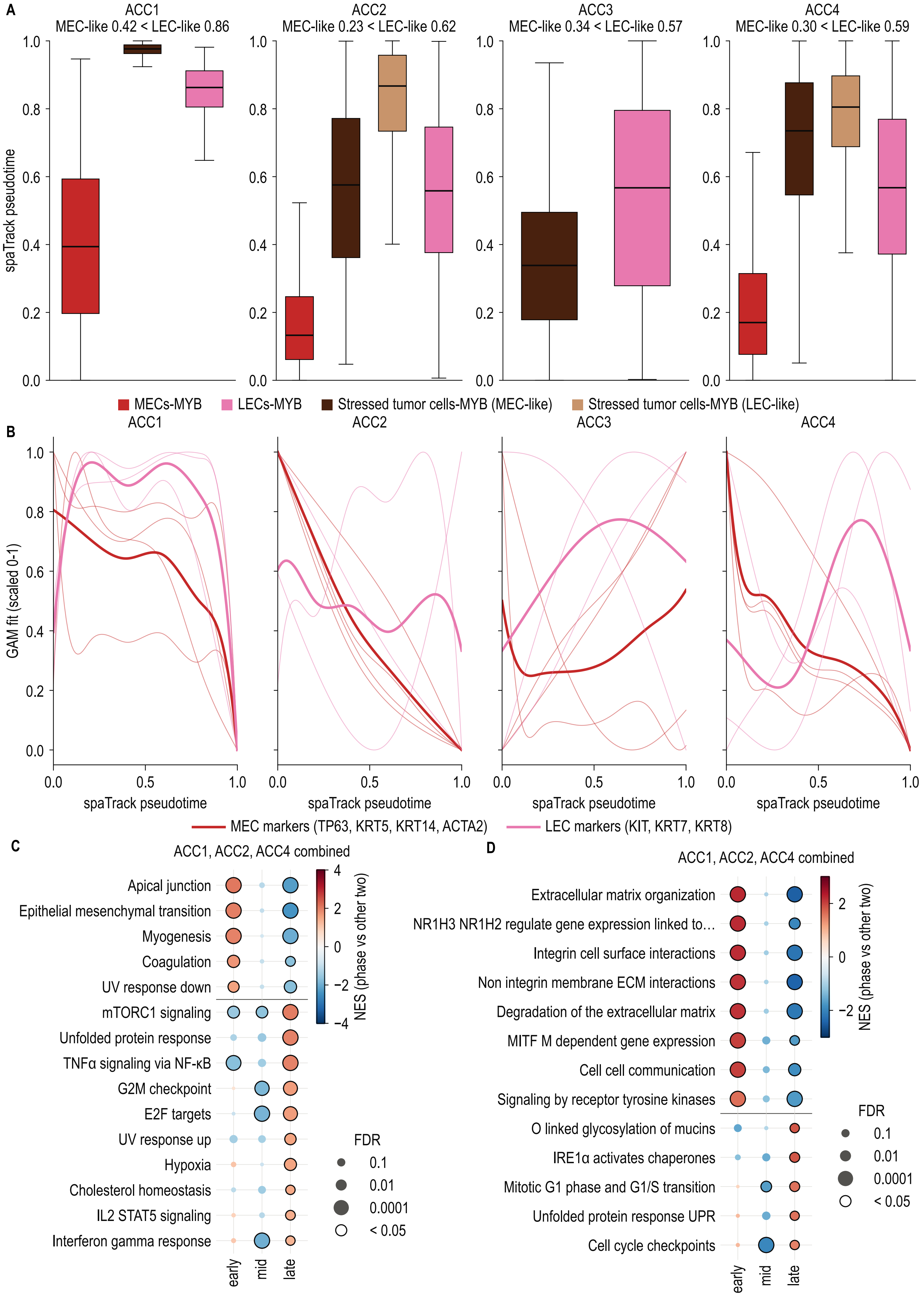

**Figure S7. spaTrack pseudotime by tumor group, marker genes and gene programs along the trajectory. (A)** spaTrack pseudotime of MECs-MYB, LECs-MYB and the MEC-like and LEC-like stressed tumor cells-MYB in each section. The median pseudotime of MEC-like and LEC-like tiles is given above each plot. Boxes as shown in Figure 5A. Shows lower pseudotime (placed earlier) for MECs-MYB compared to LECs-MYB. **(B)** MEC markers (TP63, KRT5, KRT14, ACTA2) and LEC markers (KIT, KRT7, KRT8) along spaTrack pseudotime (generalized additive model fits, scaled 0 to 1); thin lines, single genes; thick lines, mean. LEC markers rise relative to MEC markers in the later third of pseudotime in ACC1, ACC2 and ACC4; ACC3, rooted at MEC-like stressed tumor cells-MYB, does not follow this pattern clearly. **(C)** Hallmark and **(D)** Reactome gene sets enriched in the early, middle and late thirds of pseudotime. Raw counts of the tumor tiles were summed for each third and section, and each third was compared with the other two in ACC1, ACC2 and ACC4 combined (DESeq2 with section as replicate, followed by fgsea). Only genes expressed by tumor cells in the single-cell reference were ranked. (C) shows every Hallmark set with $FDR < 0.05$ and (D) the top eight Reactome sets per third. Color gives NES, dot size FDR, and a black ring $FDR < 0.05$.

Supplementary Figure Panel 8

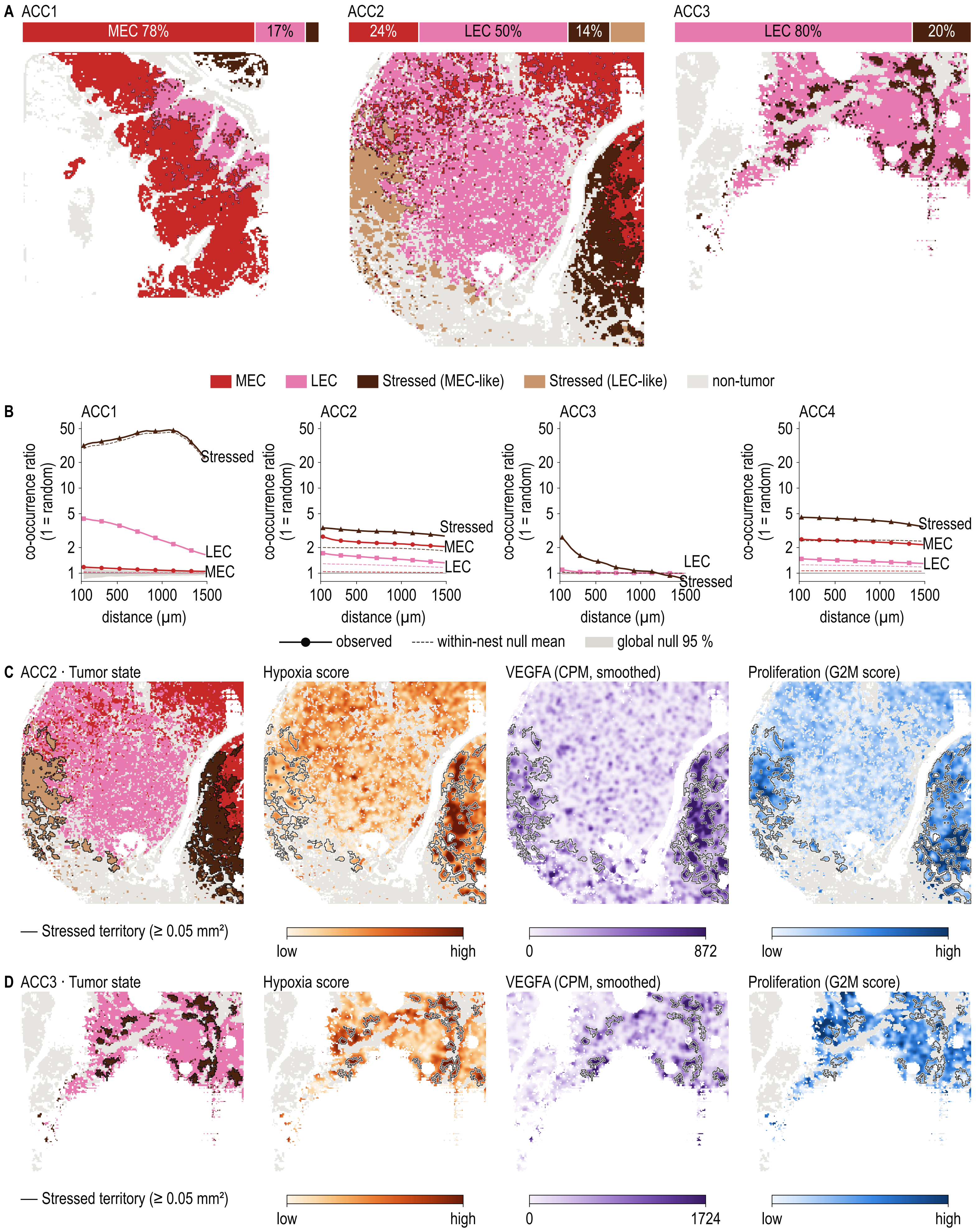

**Figure S8. Tumor-state territories and stressed-territory programs in the other sections. (A)** Tumor tiles of ACC1, ACC2 and ACC3 colored by state, with territory borders and the share of each state, on the same scale as Figure 6A. **(B)** Co-occurrence ratio against distance in each section (log scale). The ratio is the probability that a tile at a given distance shares the state of the center tile divided by the overall frequency of that state, so 1 indicates random mixing. Dashed lines show mean of label shuffles within each tumor nest; grey band (small at bottom) highlights 95% range of 1,000 global label shuffles. **(C)** ACC2 and **(D)** ACC3 maps of tumor state, hypoxia score, VEGFA CPM and G2M score, shown as in Figure 6E.

Supplementary Figure Panel 9

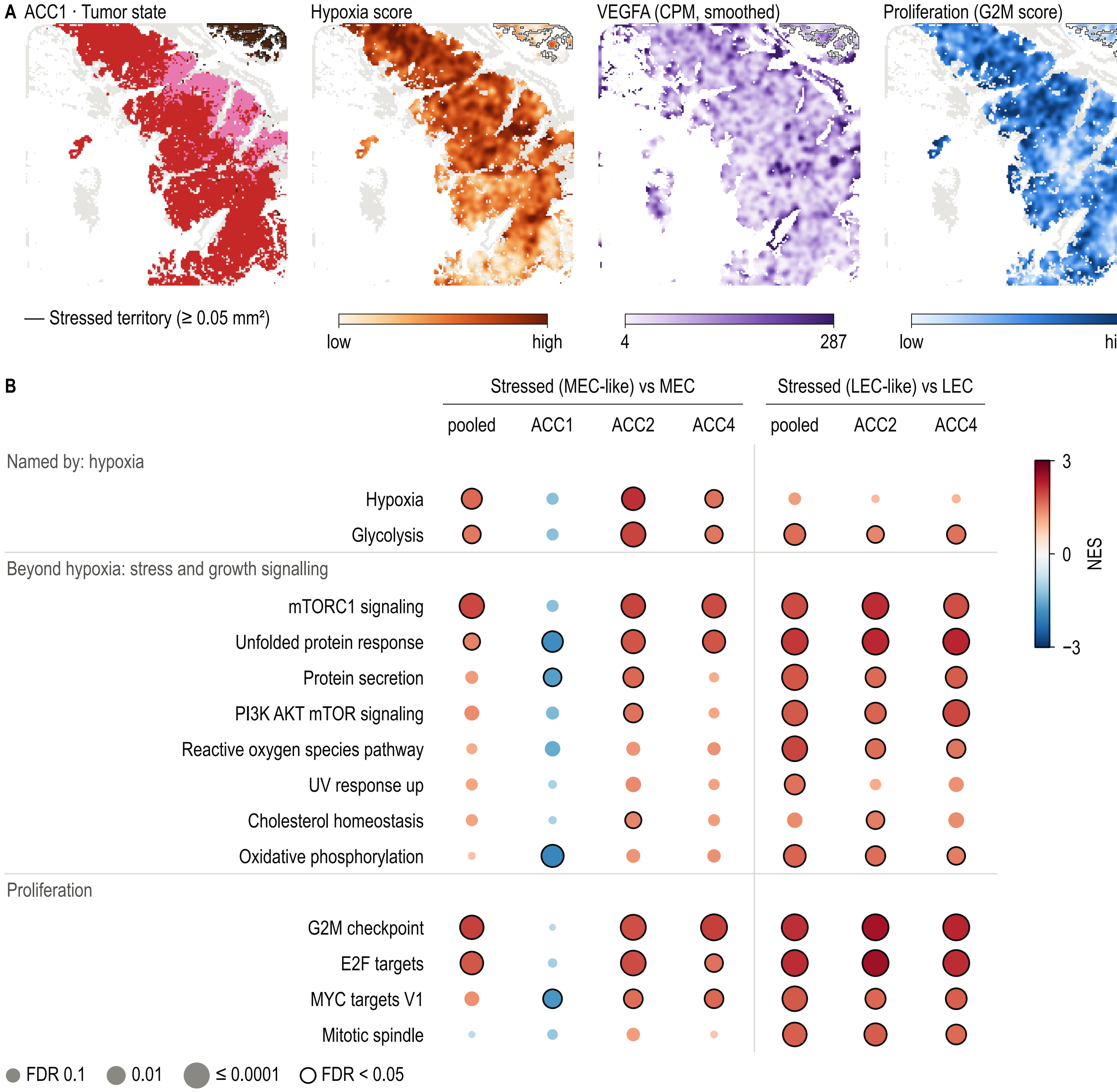

**Figure S9. Stressed-territory programs in ACC1 and comparison of each stressed subcluster with its own lineage. (A)** ACC1 maps of tumor state, hypoxia score, VEGFA CPM and G2M score, shown as in Figure 6E. **(B)** Hallmark GSEA of MEC-like stressed tumor cells-MYB against MECs-MYB (pooled, ACC1, ACC2, ACC4) and of LEC-like stressed tumor cells-MYB against LECs-MYB (pooled, ACC2, ACC4), shown as in Figure 6F. LEC-like stressed cells occur only in ACC2 and ACC4, so the pooled LEC-like comparison rests on two sections.

### Supplementary Figure Panel 10

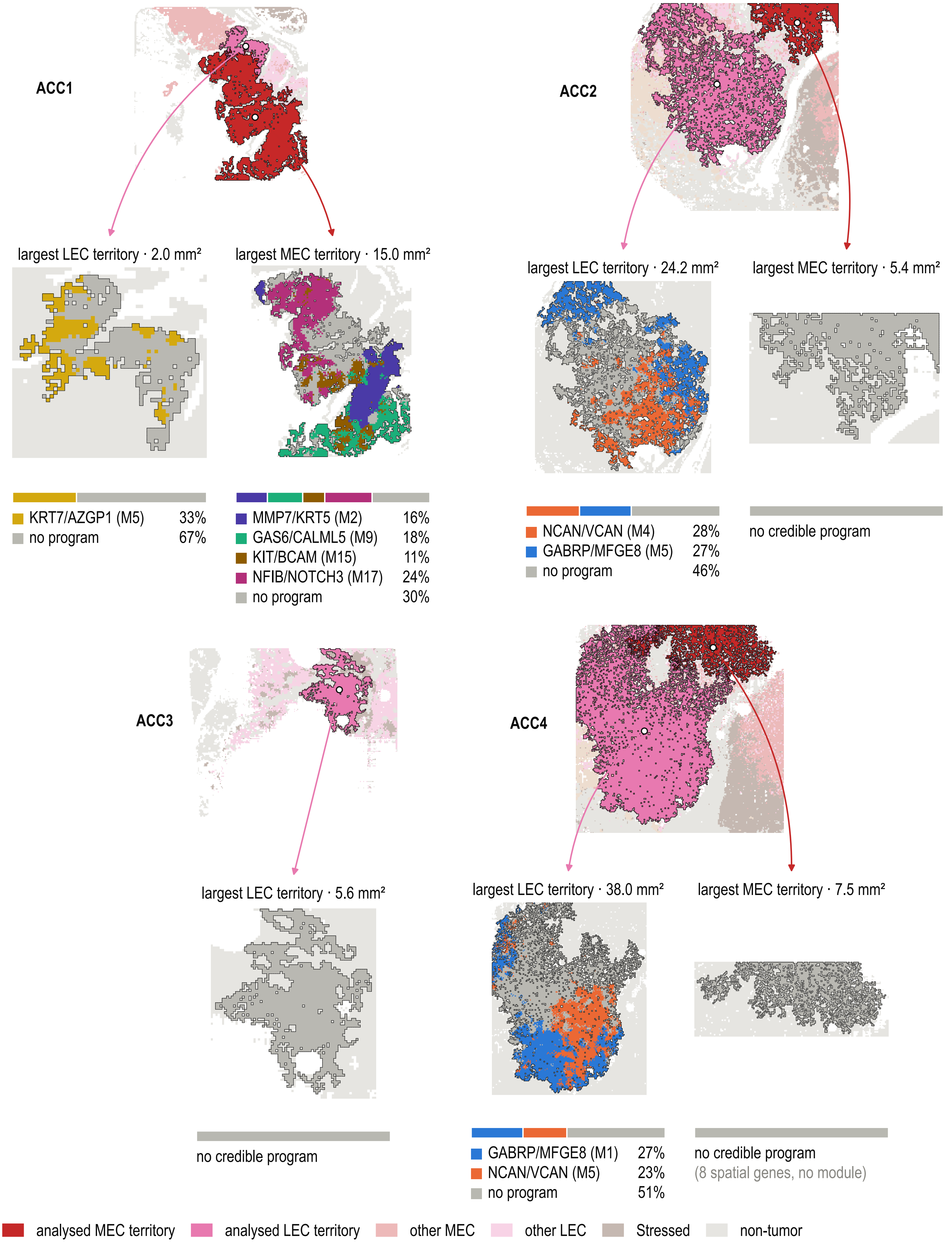

**Figure S10. Hotspot programs in the largest MEC and LEC territory of each section.** For each section, the whole tissue (top) is shown with its largest MEC and LEC territories, and arrows lead to each territory enlarged below. Bars highlight the share of bins occupied by each program identified by hotspot within the territory. Gene co-expression modules are named with two representative genes. Territory areas are given in $mm^2$.

### Supplementary Figure Panel 11

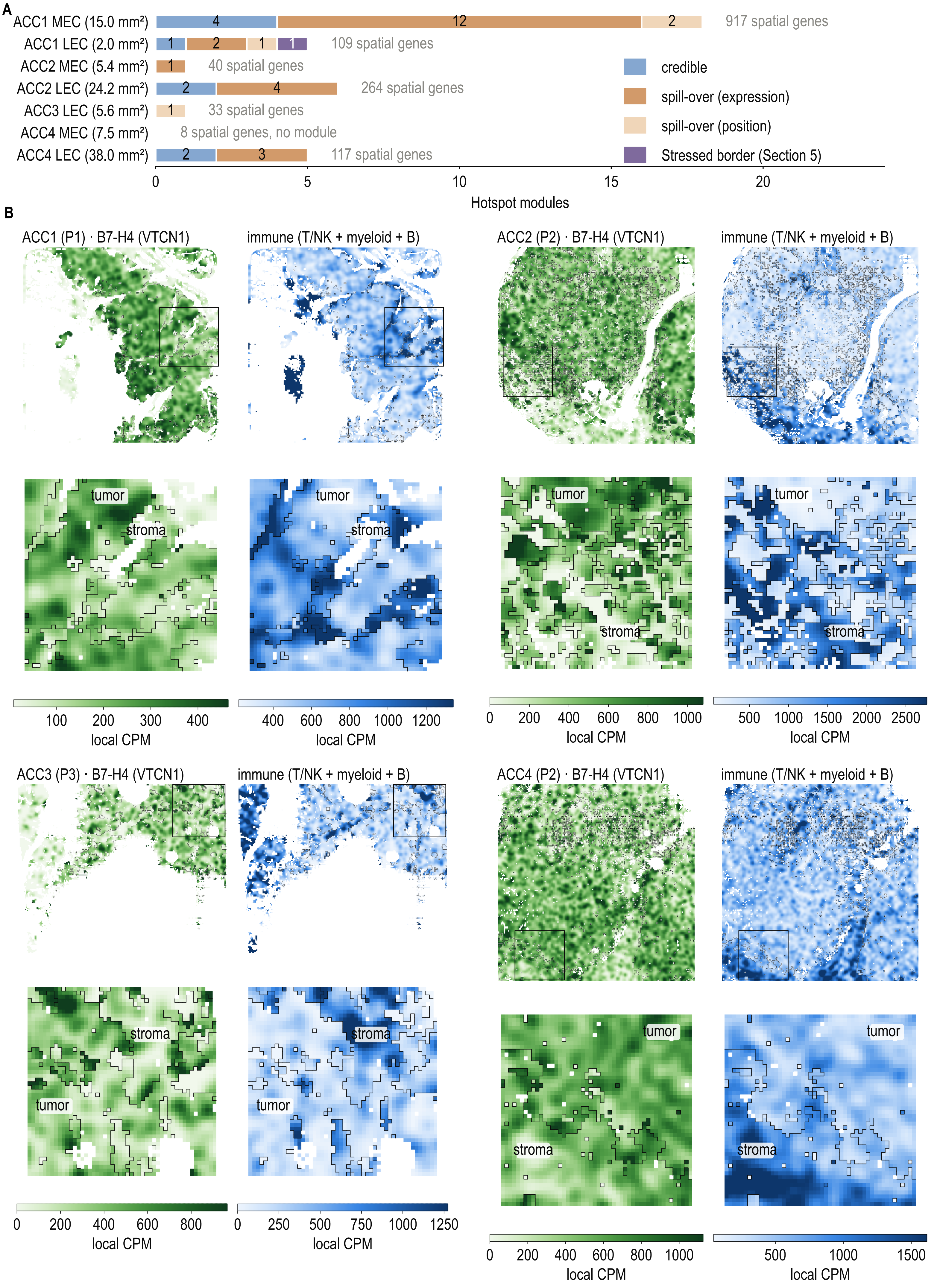

**Figure S11. Outcome of the Hotspot modules and B7-H4 against immune signal at the nest edge. (A)** Number of Hotspot modules in each territory by outcome: credible program; dropped as spill-over by expression (module genes not higher in the territory than in a neighboring compartment); dropped as spill-over by position (score rising toward a neighboring compartment); or dropped as part of the stressed-border signal. The number of spatially autocorrelated genes (FDR < 0.01) in each territory is given to the right of its bar. **(B)** Local B7-H4 (VTCN1) CPM and immune CPM (T/NK, myeloid and B-cell gene sets) in each section, smoothed with a Gaussian kernel ($\sigma$ = 75 µm) within the tumor and non-tumor sides separately. Boxes mark the 2.5 mm windows enlarged below, in which the tumor-stroma border is outlined.

**Supplementary Figure Panel 12**

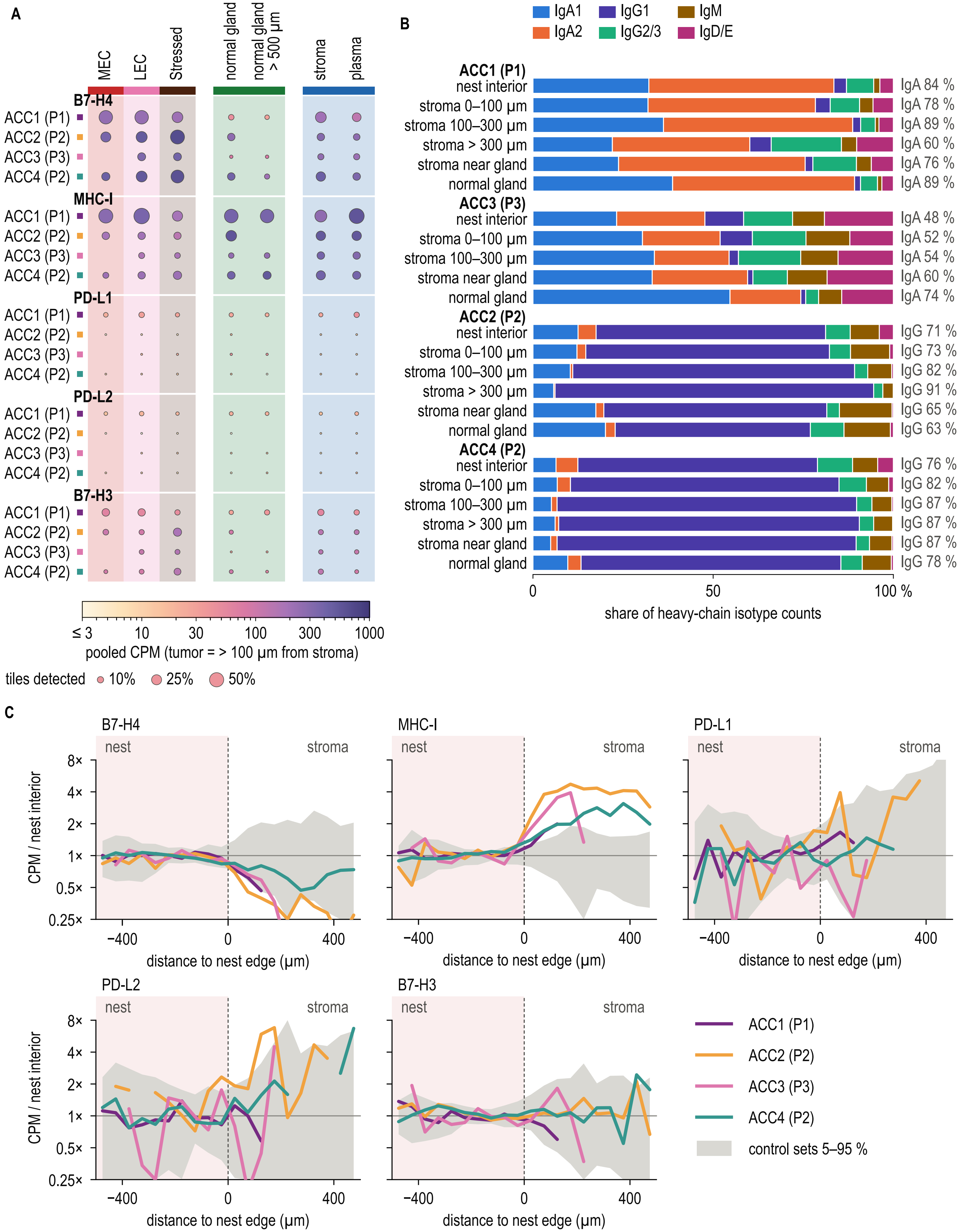

**Figure S12. Tumor-intrinsic immune-evasion genes and plasma-cell isotypes. (A)** Pooled CPM (color) and percentage of tiles with detected expression (dot size) of B7-H4 (VTCN1), MHC-I (B2M, TAP1, PSMB8, PSMB9), PD-L1 (CD274), PD-L2 (PDCD1LG2) and B7-H3 (CD276) in MECs-MYB, LECs-MYB and stressed tumor cells-MYB (tiles more than 100 µm from stroma), normal gland (all tiles and tiles more than 500 µm from tumor), stroma and plasma-cell tiles. **(B)** Share of heavy-chain isotype counts (IgA1, IgA2, IgG1, IgG2/3, IgM, IgD/E) in each zone of each section, grouped by patient: nest interior; stroma 0 to 100, 100 to 300 and more than 300 µm from tumor; stroma within 200 µm of normal gland; and normal gland. The dominant class and its share are given on the right. Zones with fewer than 50 tiles are not shown. **(C)** Tumor-intrinsic gene sets against signed distance to the nest edge, shown as in Figure 7C. P1 to P3, patients; ACC2 and ACC4 come from the same patient (P2).
